# Plant-induced transcriptional plasticity diverges between generalist and specialist herbivores

**DOI:** 10.1101/2024.12.07.627354

**Authors:** Jun Wu, Zhimou Lin, Guomeng Li, Wenyuan Yu, Yangzhi Zhang, Yishuo Kou, Pengshuai Peng, Tingfen He, Yu Wang, Shuai Zhan, Jean-Christophe Simon, Saskia A. Hogenhout, Yazhou Chen

## Abstract

Most herbivores are specialized on particular host plants but some are generalists that can exploit distinct hosts. Generalists may have evolved adaptive transcriptional plasticity to cope with the defenses of the different hosts. However, the fundamental differences in plant-induced transcriptional plasticity between generalists and specialists remain poorly understood. Here, we investigated transcriptional plasticity of the generalist aphid *Myzus persicae* and two specialist aphids, *Brevicoryne brassicae* and *Rhopalosiphum padi*, by transferring them between *Brassica napus* (a host for *B. brassicae* but not for *R. padi*) and *Zea mays* (a host for *R. padi* but not for *B. brassicae*), both suitable hosts for *M. persicae*. Generalist and specialist aphids exhibited transcriptional plasticity coordinately in response to different plant species, but their gene expression patterns often diverged. Generalists suppressed plant-defense salicylic acid (SA) signaling in the host plants, while specialist aphids provoked it in nonhost plants. SA signaling had limited effects on gene expression in the generalist aphids, but significantly shaped transcriptional responses of the specialists. These findings underscore the fundamental differences in plant-induced transcriptional plasticity between generalists and specialists and highlight the critical role of plasticity directionality in insect adaptation.

**Significance statement:** Most insects specialize in feeding on just a few specific plants, but some generalists can thrive on many plant species. This study compares the gene expression of green peach aphid *Myzus persicae*, a generalist, with two specialist aphids in response to different types of plants. Generalists showed adaptive plasticity in their gene responses, while specialists often exhibited opposite gene expression changes. Interestingly, generalists appeared to suppress plant defense signals like salicylic acid, while specialists tended to trigger them. These findings highlight how gene expression plasticity enables generalists to adapt and survive, providing new insights into the evolution of insect-plant interactions.

## Introduction

Herbivorous insects are the most species-rich among metazoans. The high diversification rate in herbivores is thought to be driven by the intimate evolutionary history of insects with their host plants (1). This is reflected in the fact that the majority of herbivores are specialized on a few closely related plant species (1). However, some insect species are extreme generalists, possessing the ability to colonize and exploit a wide variety of plant species (1). The evolutionary drivers and molecular determinants of the success of these generalist species remain largely unknown.

Aphids are excellent models to investigate the mechanisms underlying host specialization and generalism. Aphids are sap-feeding hemipterans whose stylets penetrate the plant mesophyll and navigate to the phloem for long-term feeding. Plants do not only provide food to aphids but also ecological niches that can serve several functions, such as mating, oviposition, and shelter (2). Such intensive interactions with host plants are fundamental forces driving aphid diversity (3, 4). More than 4700 species so far have been identified and the vast majority (99%) are highly specialized to one plant species or closely related ones (2). In some specialist species, it has been shown that diversification has likely occurred through divergent selective pressures from particular hosts, leading to host races that are intermediate stages of speciation, and then to true species (3, 4). Specialization can even be observed at the species level. For example, host-specialized biotypes are found in many aphid species that can only thrive on a single plant species, not even on close relatives (3, 5). Coevolution is usually invoked as a process leading to diversification through an arms race between plant defenses and insect counteradaptations. However, a shift to a different plant genus or family is often observed in closely related aphid species or biotypes, suggesting that this may be a more common route to divergence in this group of insects (8).

Among the less than 1% of species in Aphididae that are thought to be generalist species (4, 6), some have been actually found to be host-adapted biotypes or cryptic species complexes (5, 7, 9). In these cases, the species as a whole is considered as polyphagous, but individuals or populations are not. Genuine generalist species that can successfully shift from one plant to unrelated ones are therefore very rare (10). Since clonal reproduction is the main breeding system of aphids, the mechanisms underlying generalism in these aphids likely rely on gene expression changes and not on genetic variation *per se*.

The green peach aphid (GPA), *Myzus persicae*, is one of the few genuine generalists, which colonises more than a thousand plant species from over 50 plant families (2). Unlike those generalists at the species level, clonally reproducing GPA colonies that are derived from a single female can be successfully established on different plants from divergent families (11). Geographic clones of GPA collected from 14 different plant species worldwide can colonize *Brassica napus* (12). Moreover, genetically identical GPA that are reared on one plant species and then shifted to distinct plant species can survive, reproduce and form stable colonies on the new host (10, 11, 13). These findings support the idea that generalism is an intrinsic characteristic of GPA. Previous studies have shown that GPA adjustment to divergent plant hosts is associated with rapid transcriptional plasticity and involves coordinated expression of specific aphid genes in a host-specific manner (10, 11). Moreover, the upregulation of some genes in GPA is directly linked to the fitness of insects on the specific host (10, 11). This suggests that rapid transcriptional plasticity plays a crucial role in the ability of GPA to colonize diverse plant species and may therefore be the key mechanism driving its generalist ability.

Plasticity is considered adaptive when phenotypic changes align with the direction of natural selection in a given environment (14–16). Conversely, plasticity is non-adaptive when phenotypic alterations deviate from the local fitness optimum, leading to a negative correlation between the direction of plasticity and selection (14–16). Both specialist and generalist aphids exhibit the capacity for rapid transcriptional plasticity in response to different plant species (10, 17). In generalist aphids, transcriptional plasticity facilitates adjustment on diverse plant hosts, which is considered adaptive despite incurring fitness trade-offs (10, 13). In contrast, in specialist aphids, transcriptomic changes in response to nonhost plants are often non-adaptive, frequently leading to failure of colonisation (18, 19). The molecular basis underlying differences in transcriptional plasticity between generalist and specialist aphids remains largely unknown. Aphid feeding often elicits plant defense responses, particularly those mediated by jasmonic acid (JA) and salicylic acid (SA), which are key drivers of transcriptional responses in herbivorous insects (4). It remains to be determined whether plant defenses contribute to the differences in transcriptional plasticity between generalist and specialist aphids.

To investigate these questions, we used clonal lines of GPA established on the host plants *B. napus* (Bn) and *Zea mays* (Zm) derived from a single asexual female. We also included two specialist aphids *Brevicoryne brassicae* and *Rhopalosiphum padi*, known as cabbage aphid (CA) and bird cherry-oat aphid (BOA), which mainly feed on Brassicaceae and Poaceae families, respectively. We analyzed the transcriptomic changes of aphids in response to different plants and clarified the fundamental molecular differences in transcriptional plasticity between the generalist and the two specialist aphids. Furthermore, we assessed the impacts of aphid feeding on plant defenses and also investigated the roles of plant JA and SA signaling pathways in driving the divergence of transcriptional plasticity in the generalist and specialist aphids.

## Results

### Transcriptional plasticity in a generalist aphid GPA is associated with the host plants

The GPA lines that established on the host plants Bn and Zm were derived from a single female originating from a colony reared on Chinese cabbage. The lines were maintained on the corresponding plants (Bn and Zm) for more than 100 generations. GPA was more fecund on Bn than on Zm, indicating Bn plants are better hosts (*SI Appendix,* Fig. S1). To assess plasticity in the lines of GPA established respectively on Bn and Zm, we carried out plant change experiments between these two plant species by parallelly transferring aphids from the start plants to the home (same as the start plant) or the away plants (different as the start plant) (Fig. 1 *A* and *E*).

**Fig. 1.**
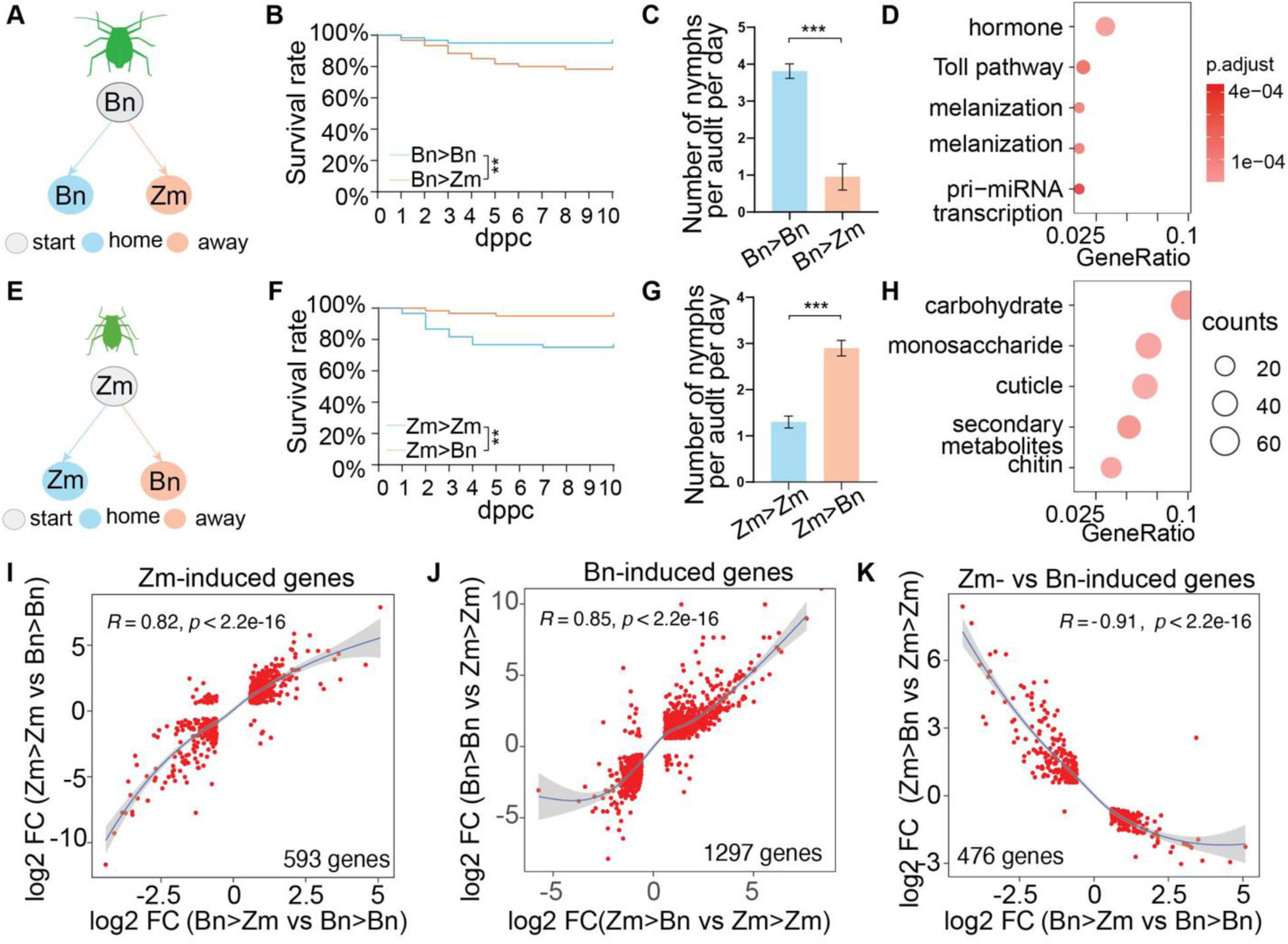
Transcriptional plasticity induced by plant changes in the generalist GPA. **A,E** Schematic overview of the experimental setup. The grey circles represent the start plants, and the blue and yellow circles represent the home and away plants, respectively. Bn is *B. napus* and Zm is *Z. mays*. **B,F** Survival rate of aphids after plant changes. The number of surviving aphids was counted each day after the plant change. ** *p*<0.01 by log-rank test. **C,G** Fecundity of aphids on home and away plants. *** *p*<0.001 by unpaired *t* test. **D,H** GO enrichment analysis for Zm-(**D**) and Bn-induced (**H**) DE genes. GO terms were detailed in Table S1 and S2. **I,J** Correlation of plant-induced gene expression changes with the basal differential expression. DE genes overlapped between basal DE and plant-induced DE genes were used for the analysis. 593 genes and 1297 genes in Zm-(**I**) and Bn-induced (**J**) DE genes overlapped the basal DE genes, respectively. **K** Correlation between Zm- and Bn-induced gene expression changes. R is Spearman rank correlation coefficient; FC is fold-change.

When GPA on Bn (GPA-Bn) were transferred, 95% of GPA remained alive on the home plants (Bn) and about 78% of them were alive on the away plants 10 days post plant change (dppc, Fig. 1*B*). When GPA on Zm (GPA-Zm) were transferred, about 75% of GPA were alive on the home plants and 95% on away plants at 10 dppc (Fig. 1*F*). The fecundity of aphids on the home plants was significantly different to that on the away plants, with GPA on Bn plants producing more offspring than on Zm plants (Fig. 1 *C* and *G*). While it is remarkable that a GPA colony originally established on Bn can effectively colonize and quickly adapt to Zm, GPA consistently performed better on Bn than on Zm.

To characterize gene expression changes induced by plant change, we performed RNA-seq analysis on survived GPA aphids on home and away plants at 2 dppc. PCA of overall gene expression revealed distinct clusters among different aphid groups (*SI Appendix,* Fig. S2*A*). Differential gene (DE) expression analysis between aphids in the home conditions (Bn>Bn vs Zm>Zm) identified 3822 DE genes (Dataset S1). These genes are referred to as basal DE genes, reflecting the basic difference of gene expression in GPA continuously influenced by the host plants Bn and Zm. Comparisons between aphids experiencing home and away conditions (Bn>Zm vs Bn>Bn, Zm>Bn vs Zm>Zm) identified 840 Zm-induced genes (Bn>Zm vs Bn>Bn) and 1857 Bn-induced genes (Zm>Bn vs Zm>Zm) (Dataset S1). Zm- and Bn-induced genes were mostly enriched for genes involved in insect hormone metabolic and carbohydrate metabolic processes, respectively (Fig. 1 *D* and *H*). Approximately, 70% of plant-specific induced genes overlapped with basal DE genes (*SI Appendix,* Fig. S2*B*), and expression patterns of these plant-specific induced genes were positively correlated with those of basal DE genes (Fig. 1 *I* and *J*). A strong negative correlation was observed between the response patterns of Zm-induced and Bn-induced genes (Fig. 1*K*). Co-expression of TPM of these genes showed coordinated up- and down-expression patterns between plant-specific induced genes and basal DE genes (*SI Appendix,* Fig. S3). These results indicate that the responsiveness of plant-induced genes in GPA is plastic and is mainly shaped by the host plants.

### Transcriptional plasticity in specialist aphids CA and BOA in response to nonhost plants

CA and BOA are specialist aphids with a limited host range, feeding mainly on Brassicaceae and Poaceae plants, respectively. To investigate the host preferences of these aphids, we conducted plant change experiments as described for GPA above (Fig. 2 *A* and *E*). At 6 dppc, 90% of CA survived on host plants (Bn), only 57% survived on nonhost plants (Zm) at 2 dppc, with none at 6 dppc (Fig. 2*B*). Similarly, 92% of BOA survived on host plants (Zm) at 6 dppc, while none of them survived on the nonhost plant (Bn) at 6 dppc (Fig. 2*F*). Offspring were produced on host plants for both CA and BOA but not on the nonhost plants (Fig. 2 *C* and *G*), confirming the high level of host specialization in these aphids.

**Fig. 2.**
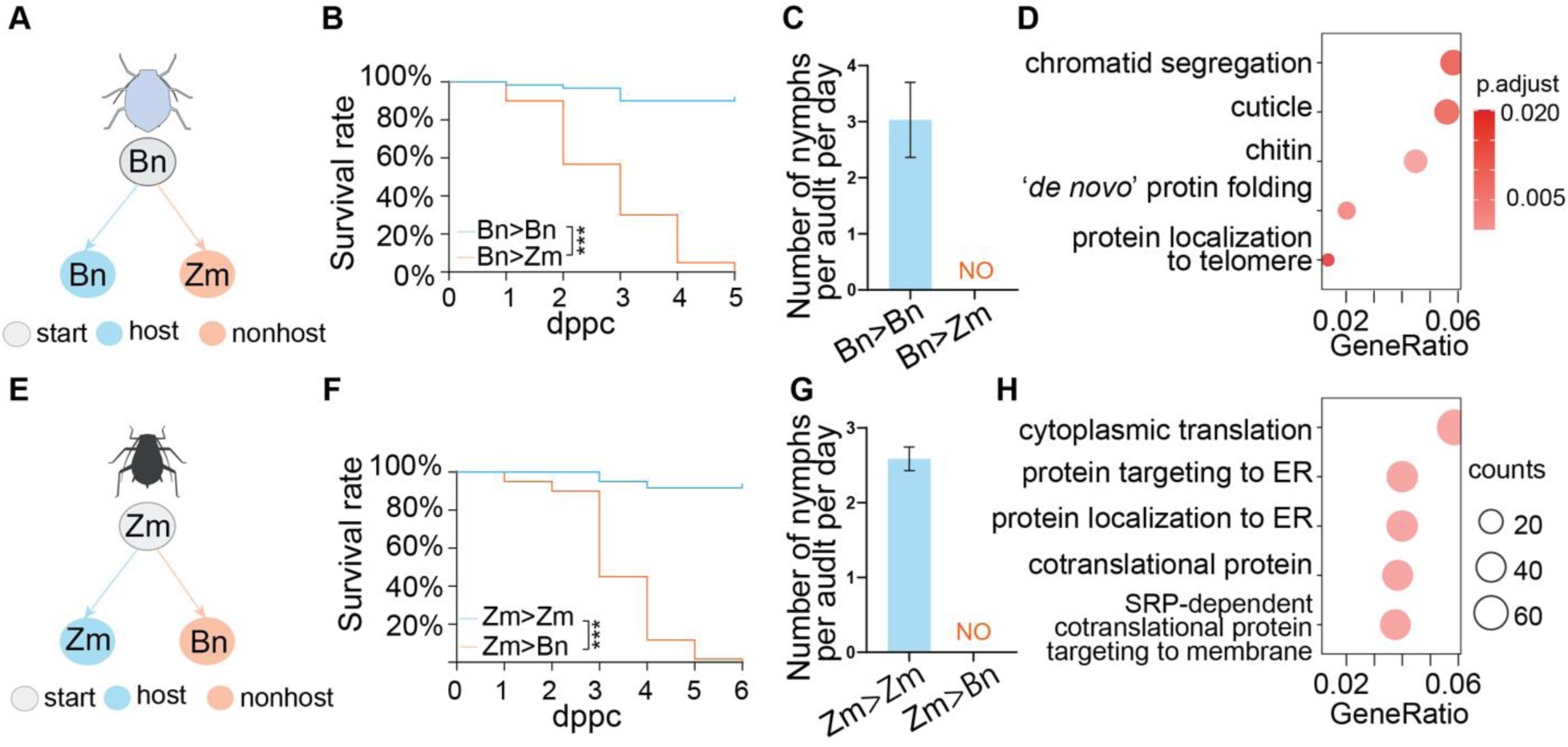
Transcriptional plasticity induced by plant changes in specialist aphids. **A,E** Schematic overview of the experimental setup. Grey circles represent the start plants, and blue and yellow circles represent the host and nonhost plants, respectively. Bn is *B. napus* and Zm is *Z. mays*. **B,F** Survival rate of aphids after plant changes. The number of surviving aphids was counted each day after the plant change. *** *p*<0.001 by log-rank test. **C,G** Fecundity of aphids on host and nonhost plants. NO means not observed. **D** GO enrichment analysis for DE genes in CA. **H** GO enrichment analysis for DE genes in BOA. GO terms were detailed in Table S3 and S4.

To analyze the transcriptional plasticity of these two specialist aphids in response to host and nonhost plants, we performed RNA-seq analysis of alive aphids on the host and nonhost plants at 2 dppc. RNA-seq reads were, respectively, mapped to the corresponding chromosome-level reference genomes (20, 21). Compared to aphids on the host plants, differential gene expression analysis identified 948 DE genes in CA and 2130 in BOA respectively induced by nonhost plants (Dataset S2). DE genes were enriched mostly in the biological processes of chromatid segregation in CA (Fig. 2*D*) and cytoplasmic translation in BOA (Fig. 2*H*). Given that aphids on nonhost plants at 2 dppc were on the path to death, we reasoned that transcriptional plasticity induced by nonhost plants in specialist aphids is mostly maladaptive.

### Plant-induced transcriptional plasticity involves genes conserved between generalist and specialist aphids

To elucidate the difference between genes associated with plant-induced transcriptional plasticity between generalist and specialist aphids, we investigated the evolutionary patterns of plant-responsive genes. We performed an all-against-all sequence comparison of protein sequences from the three aphid species, resulting in the identification of 15840 orthologous clusters. Notably, single-copy genes were significantly enriched in the DE gene sets across all four plant change experiments (Fig. 3*A*) compared to randomly selected gene sets (*SI Appendix,* Fig. S4). The enrichment of single-copy genes in the DE gene sets, irrespective of aphid species, suggests that the conserved aphid genes may play a key role in the differences in transcriptional plasticity between generalist and specialist aphids.

**Fig. 3.**
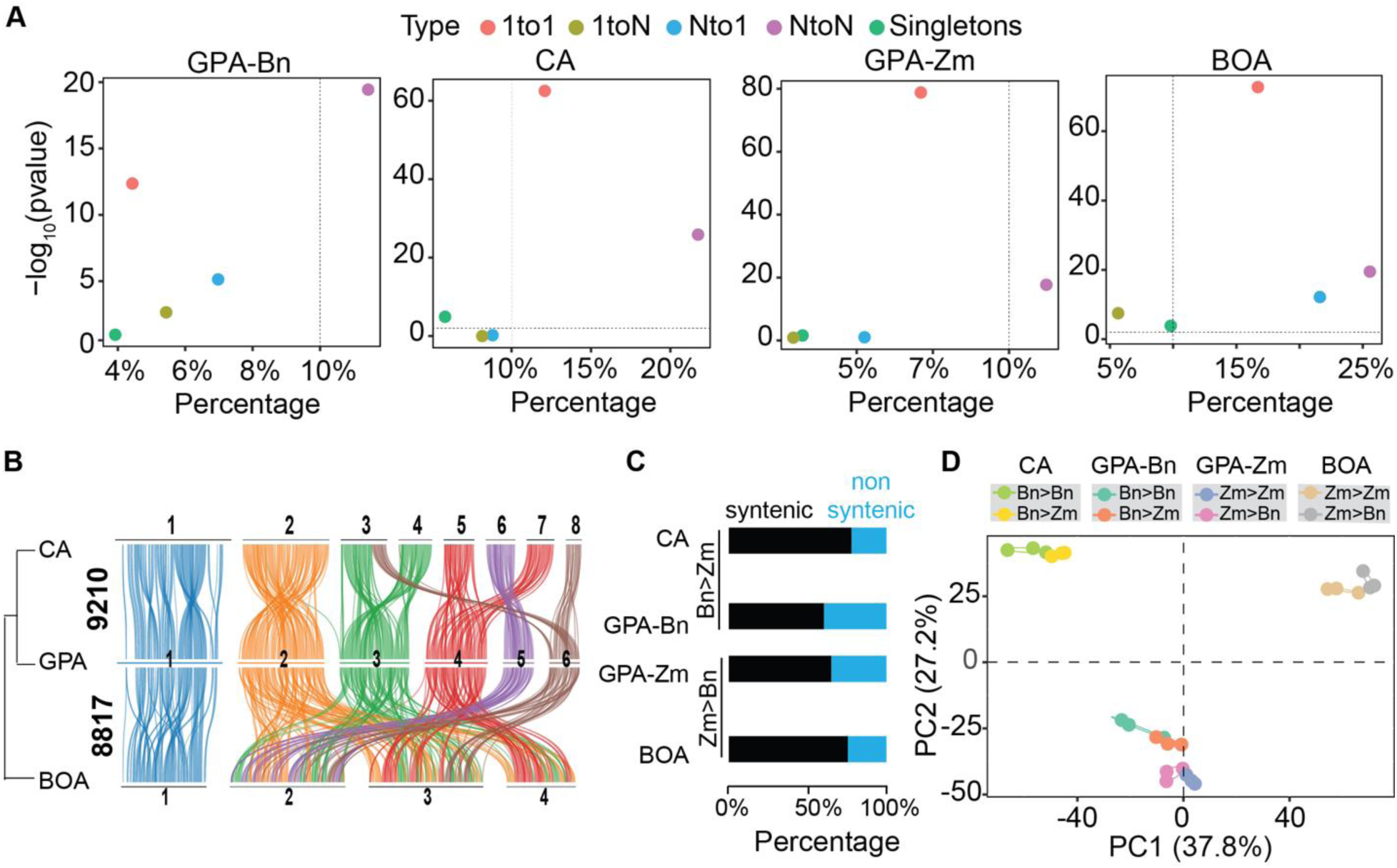
Genes involved in transcriptional plasticity enriched for conserved genes among three aphid species. **A** Enrichment of single-copy gene in the DE gene sets. To simplify the cross-species comparison, the orthologs are classified into five catalogues: single-copy genes (1-to-1), 1 to many (1-to-N), many to 1 (N-to-1), many to many (N-to-N) and singletons. The X-axis is the percentage of each category in the total DE genes. The y-axis is a log-transformed *p* value calculated by Fisher’s Exact test, estimating the probability of each catalogue in the DE gene sets compared to it in all genes in a given species. From right to left, Bn>Zm vs Zm>Zm of GPA-Bn, Bn>Zm vs Zm>Zm of CA, Bn>Zm vs Zm>Zm for GPA-Zm, Bn>Zm vs Zm>Zm for BOA. **B** Overview of syntenic blocks among three aphid genomes. The numbers with the phylogenetic tree indicate numbers of syntenic genes. The chromosomes were ordered by numbers on chromosome. Chromosome 1 is the sex chromosome. **C** Proportion of syntenic and non-syntenic genes in the DE gene sets. **D** PCA on the expression of 8459 syntenic genes.

To further characterize conserved genes between GPA and the specialist aphids, we conducted collinearity analysis, which revealed 9210 syntenic genes between GPA and CA, 8817 between GPA and BOA, and 8458 among the three species (Fig. 3*B*). Syntenic genes comprised 88.3% of single-copy genes between GPA and CA, and 88.0% between GPA and BOA (Dataset S3). The proportions of syntenic genes in the DE gene sets of CA and BOA were 74.3% and 76.4%, respectively, while the proportions in DE genes from the two host-change experiments in GPA were 58.6% and 63.5% (Fig. 3*C*).

Collinearity analysis enabled us to compare the expression patterns of homologous genes in the three aphid species that experienced the plant change. PCA on the 8459 syntenic genes across the three species revealed clear clusters of different groups. PC1 separated the three species and further distinguished aphids on the home plants from those on the away plants (Fig. 3*D*). PC2 strongly separated GPA from CA and BOA regardless of plant change (Fig. 3*D*), indicating a fundamental difference in gene expression between the generalist and specialist aphids. In addition, PC2 slightly separated the two GPA lines established on Bn and Zm. Interestingly, the responsiveness of gene expression for GPA on home versus away plants was opposite between the Bn- and Zm-lines. In contrast, the two specialist aphids exhibited consistent responsiveness of gene expression when transferred from host to nonhost plants. These data suggest the responsiveness of syntenic genes to plant changes differs significantly between the generalist and specialist aphids.

### Divergent responsiveness of conserved genes to changed plants in generalist and specialist aphids

To analyze the directionality of gene expression to plant changes, we compared the fold-change (FC) of DE genes in one aphid species to that of the syntenic ones in the counterpart species. 1041 syntenic genes between GPA-Bn and CA, and 2345 between GPA-Zm and BOA, were DE genes in at least one aphid species identified in the plant change experiments (Dataset S3). Correlation analysis on the FC of these syntenic genes showed a moderate positive correlation in gene expression between GPA and specialist aphids (*SI Appendix,* Fig. S5), suggesting the responsiveness of conserved genes diverges between generalist and specialist aphids.

To identify the genes with the most divergent responsiveness, we used a Gaussian mixture model to analyze the distribution of the FC ratio of all syntenic genes between generalist and specialist aphids (9210 between GPA and CA, 8817 between GPA and BOA). Using a probability threshold of 0.9, we identified the bias-responsive genes with changed expression greater in one aphid species compared to the counterpart species. This resulted in the identification of 404 bias-responsive genes between GPA-Bn and CA, and 503 between GPA-Zm and BOA (Fig. 4 *A* and *E* and Dataset S4). Of these, 252 genes in GPA-Bn, 152 in CA, 433 in GPA-Zm, and 70 in BOA, showed biased responses. Correlation coefficiencies for the FC of the bias-responsive genes (Fig. 4 *B* and *F*) were weak, and markedly decreased compared to that of DE syntenic genes (*SI Appendix,* Fig. S5). The overall distribution of FC of bias-responsive genes in one aphid species was significantly different from that of the syntenic copies in the other species (Fig. 4 *C* and *G*), indicating that the expression responsiveness of those genes diverged between generalist and specialist aphids.

**Fig. 4.**
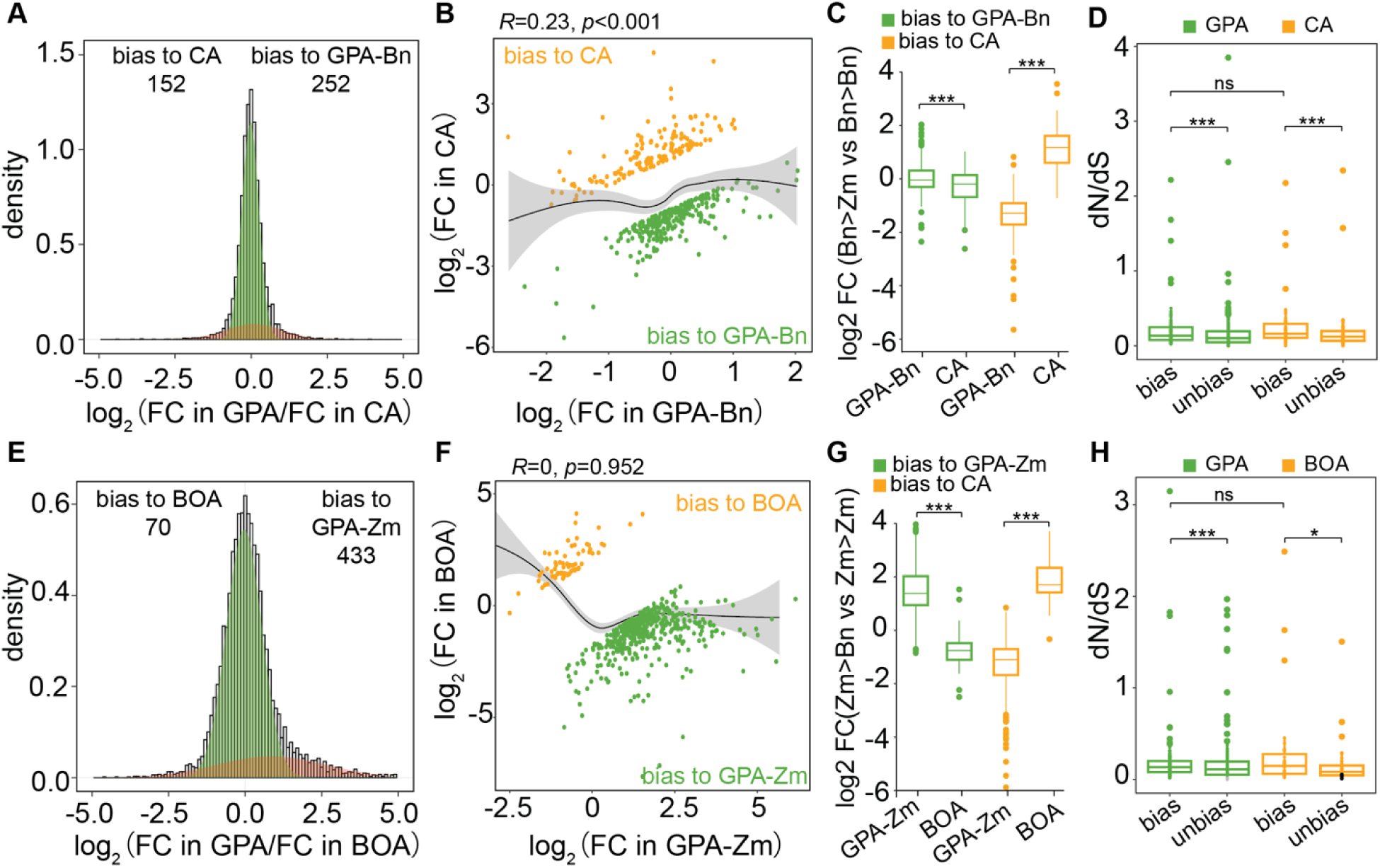
Divergent responsiveness of conserved genes in generalist and specialist aphids in response to plant changes. **A,E** Distribution of the fold-change ratio of syntenic genes between generalist and specialist aphids. The light green shading represents a Gaussian component indicating unified changes of conserved genes, and the brown shading represents biased changes in one of the aphid species. **B,F** Correlation of FC of biasedly expressed genes. Green indicates syntenic genes expressed bias toward GPA, orange indicates syntenic genes expressed bias to specialist aphids. R is Spearman rank correlation coefficient; FC is fold-change. **C,G** Comparison of FC distribution of biased expressed genes in an aphid species to the syntenic copies in the other aphid species. **D,H** Comparison of the *dN/dS* ratio of biased and unbiased expressed genes. ns is not significant, * *p*<0.05, *** *p*<0.001 by Wilcoxon test.

Overall *dN/dS* ratios for bias-responsive genes were less than 1, suggesting these genes underwent purifying selection in all three aphid species (Fig. 4 *D* and *H*). However, the ratios were significantly higher than those of non-bias-responsive genes, suggesting a relatively relaxed purifying selection for bias-responsive genes (Fig. 4 *D* and *H*). Notably, the *dN/dS* for bias-responsive genes in GPA were similar to those in specialist aphids (Fig. 4 *D* and *H*), implying both generalist and specialist aphids experienced comparable levels of purifying selection on these genes. Functional annotation of genes with bias-responsive genes revealed many belong to multigene families (Dataset S4), such as *Cathepsin B*, *lipase*, *MFS domain-containing proteins*, *UDPG*, *Cuticle Protein* genes, *Chitin-binding protein* genes. Previously, we have shown that rapid changes in the expression of *Cathepsin B* and *Cuticle Protein* genes in GPA are crucial for successful plant colonisation (10). qRT-PCR validation on aphids from plant change experiments confirmed biased expression patterns of *Cathepsin B* genes (*SI Appendix,* Fig. S6).

In summary, the responsiveness of conserved genes induced by plant change differs between the generalist and specialist aphids, possibly due to the divergent evolution of regulatory factors involved in plant interactions and host range.

### Specialist aphids provoke salicylic acid signaling of nonhost plants

Plant defenses play an important role in shaping the interactions of plants and herbivorous insects. Therefore, we switched the focus to the plant side and investigated the key plant defenses that may drive the divergence of aphid transcriptional plasticity. We first measured the plant transcriptomic responses to feeding by generalist and specialist aphids (Fig. 5 *A* and *C*) and identified plant DE genes induced by aphids compared to control plants. In Bn plants, 7309 DE genes were identified, including 630 induced by GPA-Zm and 5928 DE genes by BOA compared to control Bn plants, and 4729 DE genes between Bn plants fed by GPA-Zm and BOA (Dataset S5). In Zm plants, 997 DE genes were identified, including 288 induced by GPA-Bn, 809 DE genes by CA, and 204 DE genes between Zm plants fed by GPA-Bn and CA (Dataset S6).

**Fig. 5.**
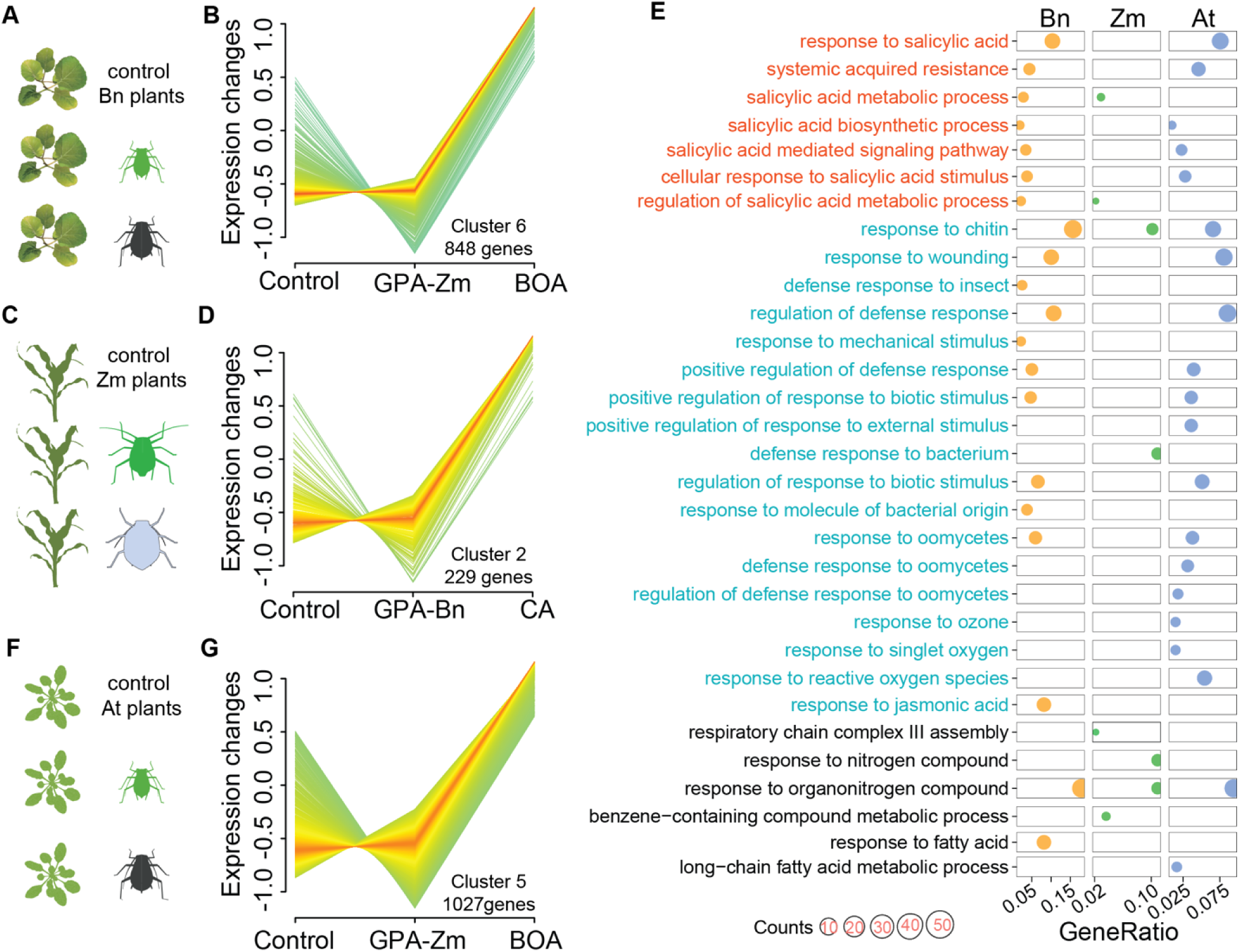
Specialist aphids provoke salicylic acid defenses in the nonhost plants. **A,C** Schematic overview of aphid feeding experiments. 40 age-synchronized 3^rd^ instar aphids were caged on the plant leaves for two days. Controls are leaves with empty cages. The caged parts of the leaves were harvested for RNA-seq analysis. Four replicates per group. **B,D** Overall gene expression pattern in the co-expression clusters. **B** Expression of 848 Bn genes in Cluster 6 in the control Bn plants, and Bn plants respectively treated by GPA-Zm and BOA. **D** Expression of 292 Zm genes in Cluster 2 in the control Zm plants, and Zm plants respectively treated by GPA-Bn and CA. **E** GO term enrichment of genes in Cluster 6(Bn), Cluster 2 (Zm), and Cluster 5 (At). Dot size represents the number of genes in the GO category. The x-axis is a log-transformed adjusted *p*-value. GO terms in red are processes related to salicylic acid pathways; GO terms in cyan are processes related to plant defense response to wounding, insect, bacterium, oomycetes, etc; GO terms in black are the other processes. **F** Schematic overview of aphid feeding experiments in At plants. **G** Expression of 1027 At genes in Cluster 5 in the control At plants, and At plants respectively treated by GPA-Zm and BOA.

To identify the plant processes that differ most caused by generalist and specialist aphids, we conducted the co-expression analysis for DE Bn genes and DE Zm genes using the TPM values. 7173 DE Bn genes were constructed into 8 co-expression clusters and 991 Zm genes were into 5 clusters (*SI Appendix,* Fig. S7 *A* and *B*). In most clusters, the responses of plant genes to specialist aphids were similar to generalist GPA. The gene expressions in those clusters were either higher or lower induced by both generalist and specialist aphids compared to control plants (*SI Appendix,* Fig. S7 *A* and *B*). Bn genes in Cluster 6 and Zm genes in Cluster 2 showed the most distinct expression patterns in response to generalists and specialists (Fig. 5 *B* and *D*). Genes in these two clusters exhibited no obvious difference in response to GPA but were dramatically induced by specialist aphids (Fig. 5 *B* and *D*). GO term analysis revealed that genes in Cluster 6 and Cluster 2 were significantly enriched for the plant pathways related to salicylic acid (SA) (Fig. 5*E*). Plant defense responses related to chitin and bacterium, that are often linked to SA pathways, were also enriched.

SA-mediated pathways are important plant defenses against herbivorous insects (23, 24). We intended to use *Arabidopsis thaliana* SA mutant plants to assess whether plant SA pathways influence aphid transcriptional plasticity. Thus, we also included *A. thaliana* wildtype (WT) in the transcriptomic analysis and compared the responses induced by GPA-Zm with those caused by BOA (Fig. 5 *F* and *G*). 4173 DE genes grouped into 5 co-expression clusters (Dataset S7 and *SI Appendix,* Fig. S7*C*). The expression patterns of genes in Cluster 5 were distinct from other clusters (*SI Appendix,* Fig. S7*C*), which exhibited no obvious difference in response to GPA-Zm but were remarkably induced by BOA (Fig. 5*G*). In Cluster 5, similar to DE genes in Bn plants, SA pathways were significantly enriched (Fig. 5*E*).

Altogether, generalists and specialist aphids differentially influence the plant SA-related defense responses. GPA is likely to alleviate the SA-related defense responses in the host plants, but specialist aphids tend to provoke them in the nonhost plants.

### Transcriptional plasticity in the generalist and specialist aphids are differentially affected by plant SA signaling

To assess the impact of plant SA signaling on aphid transcriptional plasticity, we took advantage of the available *A. thaliana* SA signaling mutant (At *npr1*) to perform plant-change experiments. We also included the JA signaling mutant (At *coi1*) in the experiments as the control to the SA signaling. GPA-Zm or BOA on the home plants (Zm) were transferred to a serial of away/nonhost plants, including Bn and three genotypes of *A. thaliana*, wild-type (At WT), At *coi1*, and At *npr1* (Fig. 6*A*). The survival rates of GPA-Zm on mutant plants were similar to those on WT plants and remained high at 8 dppc (Fig. 6 *B* and *C*). The survival rates of BOA decreased dramatically on the At WT, At *coi1*, At *npr1* plants, and no aphids survived at 8 dppc (Fig. 6 *D* and *E*). However, significantly more BOA survived on At *npr1* mutants at 7 dppc than on other *A. thaliana* genotypes (Fig. 6*E*).

**Fig. 6.**
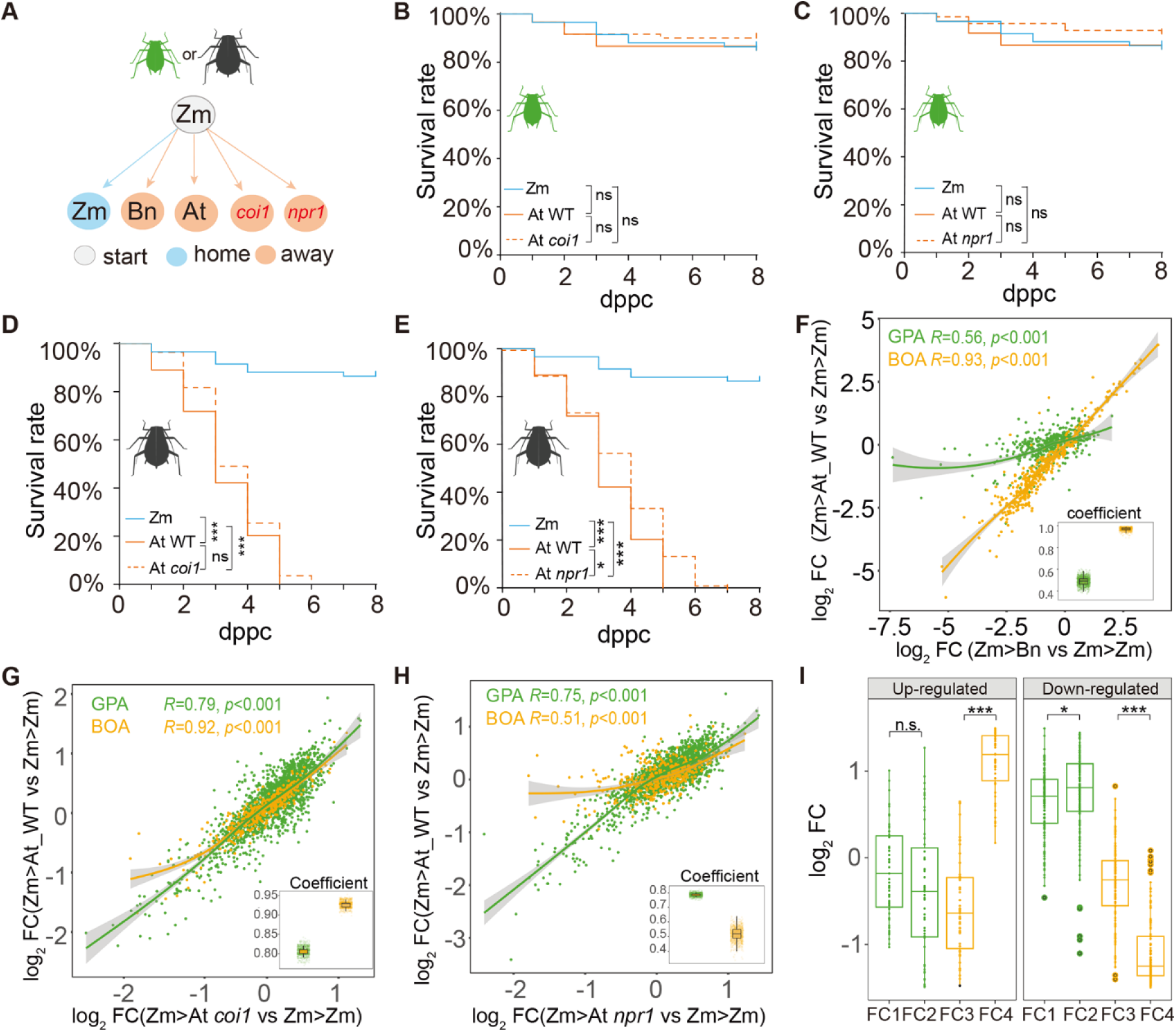
Impacts of SA and JA signaling pathways on plant-induced transcriptional plasticity in generalist and specialist aphids. **A** Schematic overview of the experimental setup. Grey circles represent the start plants, and blue and yellow circles represent the home and away plants, respectively. Bn is *B. napus* and Zm is *Z. mays*. At represents *A. thaliana* wildtype plants, *coi1* is *A. thaliana* JA signaling mutants, and *npr1* is *A. thaliana* SA signaling mutants. **B,C** Survival rates of GPA-Zm transferred from Zm to *A. thaliana coi1* (**B**) and *npr1* (**C**) mutants. **D,E** Survival rates of BOA transferred from Zm to *A. thaliana coi1* (**D**) and *npr1* (**E**) mutants. In each experiment, a total of 80 aphids were used, which were divided into 8 replicates with 10 aphids in each. The log-rank test was used to assess the significance of differences between the two survival curves. ns is not significant, * is *p* <005, *** is *p*<0.001. **F** Correlation of gene expression changes in GPA-Zm (green) and BOA (orange) induced by At plants with those induced by Bn plants. **G,H** Correlation of gene expression changes in GPA-Zm (green) and BOA (orange) induced by At plants with those induced by *coi1* mutants (**F**) and *npr1* mutants (**G**). R is Spearman rank correlation coefficient; FC is fold change. The insert plots in **F**, **G**, and **H** are bootstrapping estimated correlation coefficients. **I** FC distribution for 179 genes that biasedly and differentially expressed. Upregulated and down-regulated means BOA genes were upregulated and downregulated, respectively, when transferred from Zm to At *npr1* mutants The column heatmaps at the right display the fold changes. FC is fold change. FC1 is Zm>At vs Zm>Zm for GPA-Zm, FC2 is Zm>*npr1* vs Zm>Zm for GPA-Zm, FC3 is Zm>At vs Zm>Zm for BOA, FC4 is Zm>*npr1* vs Zm>Zm for BOA.

To analyze the aphid gene responses to wild-type and mutant plants, aphids transferred to home and away/nonhost plants at 2 dppc were subjected to RNA-seq analysis. We calculated the correlation coefficient for the FC of DE genes between the comparisons Zm>At WT vs Zm>Zm and Zm>Bn vs Zm>Zm (Dataset S8, S9). The coefficient was very high for BOA (*R*=0.93, *p*<0.001), but moderate for GPA-Zm (*R*=0.56, *p*<0.001) (Fig. 6*F*). This suggested that different nonhost plants triggered highly similar transcriptional responses in the specialist BOA, while the generalist GPA exhibited more plant-specific responses.

Using a Gaussian mixed model, we identified 1678 bias-responsive genes in GPA-Zm and 393 in BOA when transferred from Zm to At plants (*SI Appendix,* Fig. S8*A* and Dataset S10). To estimate the influence of mutant plants on aphid transcriptional plasticity, we compared the FC of these bias-responsive genes in aphids transferred to At mutants (At *coi1* and At *npr1*) with those transferred to At WT plants. For GPA, the FC of bias-responsive genes showed a strong correlation between transfers from Zm to At WT and Zm to At *coi1* (R = 0.79, *p*<0.001) or At *npr1* (R = 0.75, *p*<0.001) (Fig. 6 *G* and *H*). For BOA, the FC of bias-responsive genes remained highly correlated (R = 0.92, *p*< 0.001, Fig. 6*G*) between transfers from Zm to At WT and Zm to At *coi1*, with no significant differentially expressed (DE) genes detected under these conditions (Dataset S8). However, the correlation decreased dramatically when comparing BOA responses to At WT and At *npr1* (R=0.51, *p*<0.001, Fig. 6*H*), with 1400 DE genes detected between these two conditions (*SI Appendix,* Fig. S8*B* and Dataset S9). Overall FC distribution of BOA bias-responsive genes that also DE genes revealed that their transcriptional responses to nonhost At WT were perturbed when BOA transferred to At *npr1* from Zm (Fig. 6*I*). However, the expressions of syntenic copies in GPA were affected less by At *npr1* when transferred from Zm (Fig. 6*I*).

These results suggest that transcriptional plasticity in the generalist aphid GPA is minimally influenced by plant JA and SA signaling pathways. In contrast, transcriptional plasticity in the specialist aphid BOA to nonhost plants is influenced strongly by the SA signaling pathways but less by the JA pathways.

## Discussion

Transcriptional plasticity plays a crucial role in insects in response to environmental changes, including host plants. Generalist insects, which exploit a wide range of host plants, can rapidly modify their transcriptome according to the specific hosts, facilitating host colonisation (22, 25). In contrast, when encountering challenged hosts or nonhost plants, specialists can modify transcriptome accordingly, but this often leads to reduced fitness and even failure to colonise (18, 19). The molecular basis for the fundamental difference in the transcriptional plasticity between generalists and specialists remains largely unclear. In this study, we compared plant-induced transcriptional plasticity in the generalist aphid GPA with that in the two specialist aphids CA and BOA, by transferring aphids between Bn and Zm plants. We showed that both generalist and specialist aphids exhibit significant transcriptional changes in response to different plant species. Transcriptional plasticity is strongly associated with adaptive colonisation in the generalist aphid GPA, and with nonadaptive maladjustment in specialist aphids. Comparative genomic analysis revealed that this difference resulted from the divergent responsiveness of conserved genes between the generalist and specialist aphids, which have undergone relaxed purifying selection. GPA was likely to alleviate the plant SA signaling, but specialist aphids tend to provoke SA signaling in nonhost plants. Importantly, we found that plant SA signaling, but not JA signaling pathways, were likely involved in affecting transcriptional plasticity in specialist aphids in response to the nonhost resistance. Our findings advance previous studies by highlighting the critical roles of transcriptional plasticity in the generalist ability of GPA and emphasizing the importance of plasticity directionality in insect host colonisation.

Transcriptional plasticity enables generalist GPA to generate plant-specific transcriptomic patterns, which are strongly linked to aphid fitness on the corresponding host plants. For example, genetically identical individual GPA can colonize distantly related host species through the differential regulation of aphid genes (11, 13). GPA genes showed coordinated up- and down-regulation depending on the plant species. Knockdown of host-specific upregulated genes such as *Cathepsin B* reduced aphid fitness, but only on the hosts that induced upregulation of these genes (10). Host-specific responsive patterns of aphid genes might have evolved to avoid provoking plant defenses (26). Even when feeding on plants belonging to the same family (Bn and At), GPA exhibits dramatic differences in transcription patterns (Fig.6F). Conversely, specialist aphids like BOA show similar transcriptomic patterns when feeding on different nonhost plants, likely due to a convergent response to strong nonhost resistance, primarily mediated by the NPR1-mediated SA defense pathway. SA-mediated nonhost resistance has been reported in plants to deter pathogen infections (27, 28). Multiple lines of evidence also revealed that SA is involved in plant resistance to herbivorous insects, through a cascade of SA signaling pathways and/or crosstalk with the JA signaling (29–31). Accumulation of SA or MeSA (methyl-salicylate) induced by aphids and activation of SA-mediated plant defenses against aphid infestation have been reported in several plant species (32–36). Considering that the SA defense pathways are present in different plant species, the mechanisms of aphids to overcome the SA defense of host plants may contribute to aphid diversification through ecological divergence. Overcoming SA-mediated host/nonhost resistance may be an evolutionary driving force for novel host races.

Transcriptional plasticity has also been implicated in host adjustment in other generalists, including insects such as rice hoopers (37) and butterfly species (38), spider mites (39), and pathogens such *Sclerotinia sclerotiorum* (40). In these cases, transcriptional plasticity aligns with the necessary adjustments to the alternative host plants. This suggests a key role for adaptive transcriptional plasticity that has evolved genuine generalism. Divergent transcriptional plasticity between generalists and specialists highlighted the importance of the directionality of transcriptional plasticity. For example, when transferred from rice to wheat plants, the detoxification-related genes were upregulated in the generalist planthopper *L. striatellus*, but were downregulated in the rice specialist *N. lugens*, while ribosomal-related genes showed the opposite pattern (37). It seems that the divergent responsiveness of syntenic genes between the generalists and their closely related specialist species is a widely existing phenomenon, which may present the conserved mechanisms for herbivorous speciation.

The mechanisms underlying rapid transcriptional plasticity likely involve epigenetic regulation. Epigenetic factors, such as DNA methyltransferase (DNMT) genes and the histone modification system, have been identified in different aphid genomes (41, 42). Epigenetic mechanisms have been shown to regulate plastic traits that potentate insect adaptation (43). Moreover, expression patterns of genes often depend on their genomic landscape, such as the density of transposon elements, gene proximity, and conformation structure (44). Chromosome rearrangements between different aphid genomes may alter the landscape of conserved genes (45, 46), affecting transcriptional regulator accessibility. Indeed, extensive structural changes with many rearrangements between chromosomes have been observed between generalist GPA and the two specialist aphids (Fig. 3B), suggesting that the differences in transcriptional plasticity might be linked to chromosomal changes. Future studies profiling chromatin accessibility and quantifying the difference in syntenic genomic elements in aphid genomes with different host ranges may illuminate the genetic basis of generalism and specialization. Furthermore, RNA editing, a post-transcriptional process, plays a crucial role in transcriptional plasticity (47, 48). RNA editing alters nucleotides within RNA molecules, creating transcript diversity in response to environmental stimuli (49). These nucleotide conversions can lead to amino acid substitutions in encoded proteins, modify RNA secondary structure, create or destroy splice sites, or otherwise influence RNA processing and fate (50). These post-transcriptional modifications ultimately fine-tune gene expression patterns, contributing to transcriptional plasticity.

In summary, plant-induced transcriptional plasticity diverges between generalist and specialist aphids, which is influenced by the plants they are associated with, ultimately affecting their success or failure in colonisation.

## Materials and Methods

*M. persicae* (GPA) clone was obtained from the stock on cabbage plants (*B. oleracea*) in the lab of Prof. Tongxian Liu at Northwest A&F University. Two lines (GPA-Bn and GPA-Zm) were maintained on host plants over two years. *B. brassicae* (CA) was collected from rapeseed fields in Lanzhou and maintained on Bn in our lab. The clone of *R. padi* (BOA) was obtained from the stock on wheat plants in the lab of Prof. Julian Chen at the Chinese Academy of Agricultural Sciences and maintained on Zm in our lab. All aphid stocks were kept under controlled conditions (24 ± 1°C, 60 ± 10% RH, 16/8 h light/dark) and had been maintained in the lab for over two years. *Z. mays* (Zm) B73, *B. napus* (Bn) Zhongshuang 11, and the *A. thaliana* plants were grown under identical conditions. Arabidopsis mutants *npr1-2* were kindly provided by Prof. Huan Chen (Shanghai Jiao Tong University, China) and *coi1-2* by Prof. Yingbo Mao (CAS Center for Excellence in Molecular Plant Sciences, China).

For plant change experiments, ∼100 adult aphids from each colony (GPA-Bn, GPA-Zm, CA, and BOA) were transferred to 4-week-old host plants. After 24 hours, adults were removed, and newly born nymphs were reared to the 3^rd^ instar. Thirty age-synchronized 3^rd^ instar aphids were then transferred between home and away plants. Transfers were repeated 3–5 times to ensure at least three replicates for RNA-seq. For aphid feeding experiments, 30 3rd instar aphids were caged on 4-week-old plants for 48 hours. Plants were washed with double-distilled water to remove aphid traces, and three replicates per group were harvested for RNA-seq.

Samples were homogenized using a TissueLyser III (QIAGEN), and total RNA was extracted using TRIzol® (Invitrogen) and VAHTS® mRNA Capture Beads (Vazyme). Strand-specific RNA-seq libraries were prepared using the VAHTS® Universal V6 RNA-seq Library Prep Kit and sequenced on the Illumina NovaSeq 6000 platform at OEbiotech (Shanghai). RNA-seq analyses were performed with Read Mapping and Transcript Assembly (RMTA) (v2.6.3) and the edgeR (v3.0).

Gaussian mixture modeling of syntenic gene log-transformed fold changes was performed using the normalmixEM function (mixtools). Co-expression clusters were identified using Mfuzz (v2.6.1). All data analyses were performed in R. All statistical tests are described in the figure legends. More details on the materials and methods used in this study are provided in *SI Appendix*.

## Acknowledgments

We thank Prof. Tongxian Liu (Guizhou University, China), Prof. Julian Chen (The Institute of Plant Protection, Chinese Academy of Agricultural Sciences, China) for kindly providing the aphid colonies. We also thank Prof. Yingbo Mao (CAS Center for Excellence in Molecular Plant Sciences, China) and Prof. Huan Chen (Shanghai Jiao Tong University, China) for the *A*. *thaliana* mutants.

This project is funded by the National Key Research and Development Program of China (project No. 2023YFF1000703 to YC) and the National Natural Science Foundation of China (project No. 32172392 to YC), and supported by Hubei Hongshan Laboratory (project No. 2022hszd026 to YC), the Startup Foundation for Advanced Talents at HZAU to YC, the Fundamental Research Funds for the Central Universities (Program No. 2022ZKPY003 to YC), and the Wuhan Yingcai Talent Program to YC.

## Supporting Information

### Materials and Methods

#### Plant materials

*Z. mays* (Zm) B73, *B. napus* (Bn) Zhongshuang 11, and the *A. thaliana* plants were grown under conditions with 16 h light/8 h dark at the temperature 24 ± 1°C at day and 20 ± 1°C at night. Arabidopsis mutants *npr1-2* were kindly provided by Prof. Huan Chen (Shanghai Jiao Tong University, China) and *coi1-2* by Prof. Yingbo Mao (CAS Center for Excellence in Molecular Plant Sciences, China). The mutants *npr1-2* and *coi1-2* are point mutations, which were verified by sequencing of PCR products with previously reported primers (Table S5) (1, 2). 4-week-old seedlings were used for the experiments.

#### Insect rearing

*M. persicae* (GPA) clone was obtained from the stock on cabbage plants (*B. oleracea*) in the lab of Prof. Tongxian Liu at Northwest A&F University. GPA was initially reared on Bn for 2-3 weeks, then transferred to Zm to establish a stable colony. Two colonies (GPA-Bn and GPA-Zm) were maintained on corresponding host plants for more than two years. *B. brassicae* (cabbage aphid, CA) was collected from a rapeseed field at Lanzhou by Prof. Wenhong Li and maintained on Bn in our lab. The clone of *R. padi* (bird cherry-oat aphid, BOA) reared on wheat plants was obtained from the lab of Prof. Julian Chen at the Chinese Academy of Agricultural Sciences and maintained on Zm in our lab. All the aphid stocks were kept on the plants grown at 24 ± 1°C, 60 ± 10% relative humidity and a 16/8 h light/dark photoperiod cycle in the growth chamber. Aphid stocks used for all experiments have been maintained in the laboratory for more than two years.

#### Plant change experiments and aphid sample collection

Approximately 100 adults of each aphid colony (GPA-Bn, GPA-Zm, CA, and BOA) were collected from the stocks and then transferred to 4-week-old host plants. The next day, adults were removed from plants and newly born nymphs were kept on the plants until grew into the 3^rd^ instar. Each of the 30 age-synchronized aphids (3^rd^ instar) was transferred from the start plants parallelly to the home plants and away plants. The transfer was repeated three to five times, achieving at least three replicates. To avoid escaping, aphids were confined on plant leaves in foam clip-cages (2 cm height × 3.5 cm diameter) sealed at the base of the petiole. Two days after plant change, alive aphids from both home and away plants were collected for RNA extraction, which was subsequently utilized for RNA sequencing and quantitative reverse transcription PCR (qRT-PCR) analysis. The wingless asexual female was used in all plant change experiments.

#### Aphid survival rate assay

The survival rates of GPA-Bn, GPA-Zm, CA, and BOA on Bn and Zm plants (Fig. 1 B and F, Fig. 2 B and F) were conducted on leaf discs. Ten age-synchronized aphids (3^rd^ instar) were transferred to leaf discs of either home or away plants. Each leaf disc was prepared by placing a fresh detached leaf from home or away plants on a thin layer of water agar within small petri dishes (3.5 centimeters in diameter). Fresh leaf discs were replaced every two days. This transfer process was repeated 8 to 12 times, ensuring a minimum of 8 replicates. The number of alive aphids was counted every day. The survival rates of GPA-Zm and BOA on Zm, Bn, and At plants (Fig. 6 B and E) were conducted on the whole plants with aphids caged on leaves. Ten age-synchronized aphids (3^rd^ instar) were caged on leaves and alive aphids were counted every day. The log-rank test was used to assess the significance of differences between the survival curves of aphids on home and away plants.

#### Aphid infestation and plant sample collection

Four-week-old Bn, Zm and At plants were treated with 30 age-synchronized GPA-Zm and BOA for 48 hours, respectively. Four-week-old Zm and Bn plants were treated with 30 age-synchronized GPA-Bn and CA for 48 hours, respectively. Wingless asexual females were confined to leaves using foam clip-cages (2 cm height × 3.5 cm diameter), the controls were leaves of clear plants with empty cages. Three replicates for each group were harvested. After 48 hours of aphid feeding, the caged plant tissues were carefully washed with double-distilled water to eliminate aphid traces, then rapidly stored in 1.5 ml nuclease-free Eppendorf tubes and immediately frozen in liquid nitrogen for RNA-seq analysis.

#### RNA sequencing

The aphid and plant samples were ground using tissue homogenizer TissueLyser III (QIAGEN). The total RNAs from samples were extracted using TRIzol^®^ Reagent (Invitrogen) in combination with purification by VAHTS^®^ mRNA Capture Beads (Vazyme, Nanjing, China). The concentration and purity of RNA samples were determined using NanoDrop 2000 and Qubit 2.0 Fluorometer. The integrity of the RNAs was assessed with the Agilent 2100 Bioanalyzer. The strand-specific paired-end libraries were constructed using the VAHTS^®^ Universal V6 RNA-seq Library Prep Kit (Vazyme, Nanjing, China) following as the instruction manual, were then sequenced by the Illumina Novaseq 6000 platform at OEbiotech (Shanghai, China).

#### Genome annotation

The chromosome-level genomes of GPA, CA, and BOA were published previously (3–5). We reannotated the BOA genome with the same annotation pipeline to achieve a similar number of protein-coding genes as annotated in GPA and CA. Braker pipeline was used for protein-coding gene prediction, which combines gene models predicted based on short-read RNA-seq data (Braker1 method) (6) and protein homologs from other aphids (Braker 2 method) (7). For Braker1, transcriptome data from wingless/winged asexual females and nymphs were used. RNA-seq reads were mapped to the reference genome using HISAT2 (v2.2.1) (8) and processed by SAMtools (v0.1.18) (9). For Braker2, protein-coding genes from *M. persicae* (3), *A. pisum* (10) *A. gossypii* (11), and *Drosophila melanogaster* (12) were used as homolog evidence. The two generated gene sets were compared at both exon and transcript levels to select the best gene models. Consistent gene models and unique models non-overlapped to each other were selected first, while the models with the disparity between the two approaches were further checked with homolog alignment and transcriptome mapping results.

#### Differential expression analysis

RNA-seq analyses were performed with Read Mapping and Transcript Assembly (RMTA) app (v2.6.3) and the edgeR (multifactorial pairwise comparisons) app (v3.0). Briefly, clean reads were mapped to the reference genome with aligner HiSat2 with default settings, and then reads were quantified using the FeatureCounts. Read counts were submitted to the edgeR app with default parameters at the platform CyVerse Discovery Environment (DE CyVerse). The option Reference biological condition was set depending on experiments.

At the plant change experiments, aphid samples collected from home plants were set as reference biological conditions. Aphid genes with expression change more than 1.5-fold with an adjusted *p*-value less than 0.05 were considered as differentially expressed. For plant genes, expression changes more than 2-fold with an adjusted *p*-value less than 0.05 were considered as differentially expressed.

Gene expression levels were determined using the TPM (Transcripts Per Million) method, which was performed with the TPMCalculator (v0.0.3). The BAM files, which were produced by the RMTA (v2.6.3) process, were input into the TPMCalculator for analysis. Utilizing the annotation GTF file, TPM values were computed for each gene.

#### GO term enrichment analysis

GO term enrichment analyses using the clusterProfiler package (v3.18.1) (13). The input for these analyses was a series of defined DEG IDs. Default settings were used.

#### Ortholog analysis

Protein sequences of GPA, CA, and BOA were used for orthology analysis using the pipeline of OrthoFinder (v2.5.4) (14) with the parameters “-M msa -S blast”. This pipeline resulted in 8,067 single-copy universal genes with strict single-copy across three species and 15,840 orthologous clusters.

#### Phylogenetic analysis

Maximum likelihood (ML) phylogenetic trees for the aphid gene family were constructed using protein sequences from each gene family member across the three aphid species with RAxML software (v8) (15) with default parameters. The multiple sequence alignment was performed using ClustalW (v2.1) (16), and the resulting ClustalW-formatted FASTA files served as input for RAxML to build the ML trees. To evaluate the robustness of the tree branches, a bootstrap analysis with 1,000 replicates was conducted. The online tool iTOL (v6) (17) was utilized for visualizing and annotating the phylogenetic trees.

#### Syntenic analysis

Syntenic analysis was performed by One Step MCScanX at TBtools (18). Briefly, the genome files and annotation GFF files were used to process pairwise comparisons of collinearity and generate a serial of files (collinearity, ctl, and gff). The same type of files was merged by the plugin File Merge For MCScanx. SynVisio was used to visualize the genome synteny.

#### PCA analysis and heatmap

PCA analyses were performed using the R package ggbiplot. A matrix of normalized read counts of genes generated by edgeR was used for PCA analyses. Heatmap was generated by the R package ComplexHeatmap (19). The *Z*-score transform was based on a row.

#### Gaussian analysis

Gaussian mixture modelling was performed on log-transformed fold-change for all syntenic genes. The normalmixEM function from the R package mixtools was used for the modeling. Specifically, we restrained the number of normal distributions to two, and the two distributions were homoscedastic. For example: y = normalmixEM(x, lambda = 0.5, mu = c(0, 2), sigma = (0.5)).

#### dN/dS ratio

We calculated the rates of synonymous substitution per synonymous site (*dS*) and non-synonymous substitution per non-synonymous site (*dN*) for genes with biased expression between generalist aphids (GPA-Bn and GPA-Zm) and specialist aphids (CA and BOA). Initially, we identified orthologous proteins across the three aphid species and generated protein alignments using MUSCLE (v3.8.31) (20). These alignments guided the codon alignments of the coding sequences (CDS) for each pair of orthologous genes using PAL2NAL (21). Pairwise dN and *dS* values were then calculated from these codon alignments using PAML (v4.4) (22) with the YN00 model. We excluded orthologous gene pairs with *dS* values > 2 from our analysis to account for potential saturation effects, which can distort substitution rate estimates. To ensure the accuracy of evolutionary rate comparisons (*dN/dS*), we applied stringent filtering criteria to avoid erroneous estimates due to insufficient sequence divergence; pairs with *dN* or *dS* values < 0.01 and fewer than 50 synonymous sites were excluded.

#### Mfuzz cluster analysis

Co-expression cluster analysis was performed by the R package Mfuzz (v2.6.1) (23). Using the soft clustering method of fuzzy c-means clustering, we determined the mean gene expression values for each treatment group. These mean values were then fed into the fuzzy c-means algorithm, resulting in the identification of distinct gene expression clusters, each exhibiting unique profiles.

#### qRT-PCR analysis

Total RNA was isolated from adults using TRIzol® Reagent (Invitrogen) and subsequent DNase treatment using an RNase-free DNase I (Thermo Fisher Scientific). cDNA was synthesized from 1 μg total RNA with RevertAid First Strand cDNA Synthesis Kit (Thermo Fisher Scientific). The qRT-PCR reactions were performed on the CFX Connect™ Real-Time System (BIO-RAD) using gene-specific primers (Table S6). Each reaction was performed in a 20 μL reaction volume containing 10 μL SYBR Green (Thermo Fisher Scientific), 0.5 μL of each primer (10 mM), 1 μL of sample cDNA, and 8 μL double distilled water. The cycle programs were: 95°C for 10 s, 40 cycles at 95°C for 20 s, 60°C for 30 s. Relative quantification was calculated using the comparative 2^−△Ct^ method (24). All data were normalized to the level of the *ribosomal protein L7* (*RPL7*) and the *nuclear elongation factor 1-alpha (EF1α)* genes from the same sample.

The design of gene-specific primers was achieved by three steps. Firstly, NCBI Primer-Blast was used to obtain two to three pairs of PCR primers for each gene. Next, the primers were blasted against the cDNA database of the target aphid species to ascertain the specificity. Third, the melting curves of PCR products were examined carefully, the products were sequenced to validate their correctness. Only primers amplified correct fragments with sharp melting curves were used for qRT-PCR analyses (Table S6).

#### Statistical analysis

Correlation analysis and Fisher’s Exact test were performed in R. The survival rate of aphids was analyzed by the log-rank test. The correlation of gene expression changes was estimated by Spearman rank correlation coefficient. Bootstrapping correlation coefficients were performed using *cor.test* function from R package Tidymodels (25).

#### Data deposition

RNA-seq data of aphid samples (GPA, CA, and BOA) were deposited in the NCBI BioProject under project numbers PRJNA1138887 and PRJNA1194682, and RNA-seq data of plant samples were under PRJNA1194732. The subset of CA RNA-seq data was previously deposited under the project PRJNA1104693 with the accession numbers SRR28892653, SRR28892654, and SRR28892655.

### Supplementary Figures

**Fig. S1.**
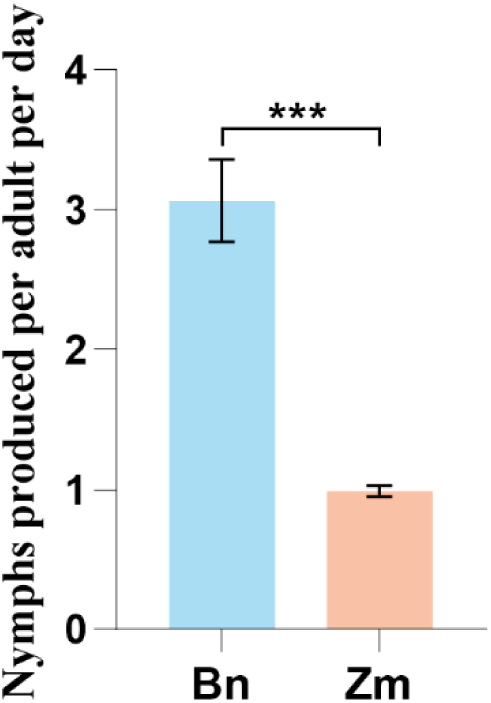
Number of nymphs produced by GPA on Bn and Zm plants. A single 1^st^ instar GPA-Bn or GPA-Zm was reared on Bn or Zm plants. When aphids developed into adults, the number of nymphs was counted every two days for a total of 10 days. Six to eight replicates in each group. Data are shown as mean ± SEM. Statistical significance was determined using the unpaired *t* test. * *p*<0.05, ** *p*<0.01 and *** *p*<0.001.

**Fig. S2.**
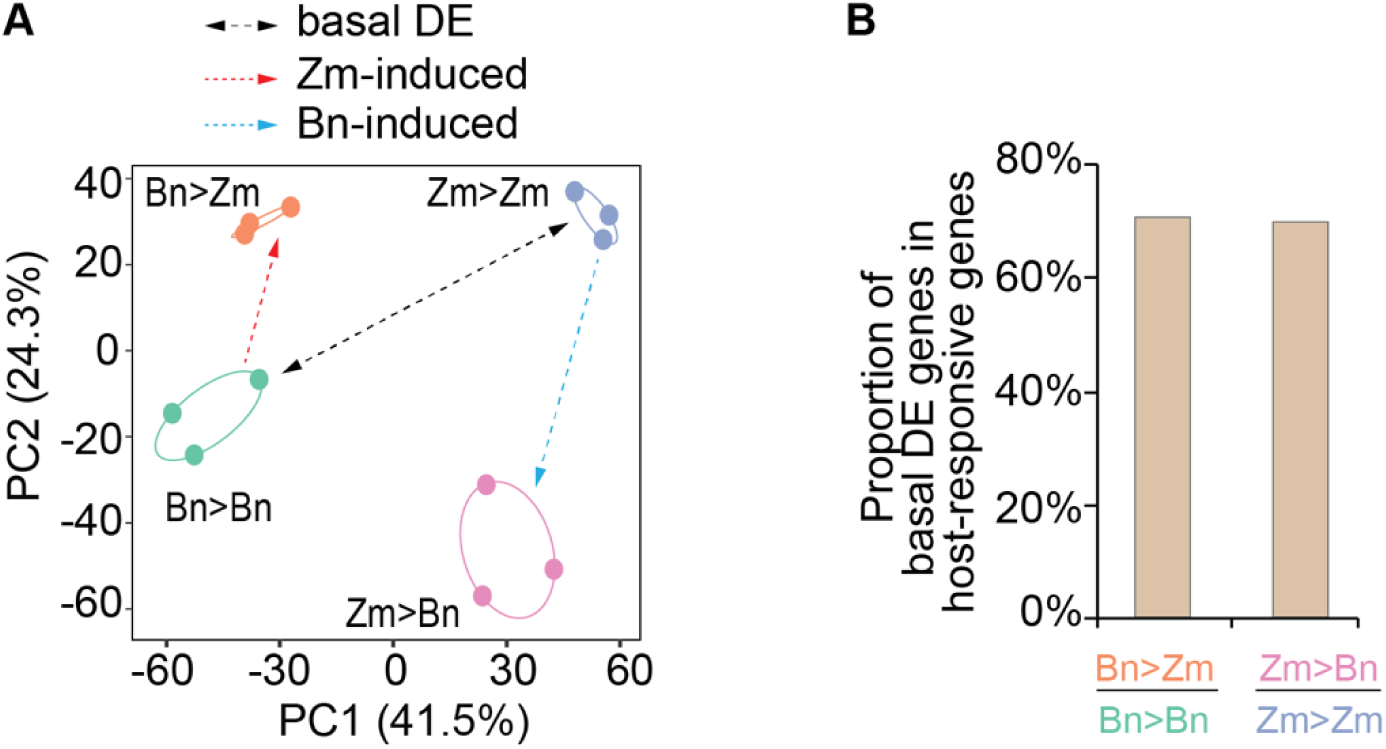
Transcriptomic analysis of GPA-Bn and GPA-Zm experienced plant changes. **A** PCA based on overall gene expression separated different sample groups. The black dashed line represents the basal difference revealed by the comparison between GPA-Bn and GPA-Zm transferred between home plants. Orange and blue dashed lines represent the Zm-induced and Bn-induced DE genes revealed by the comparisons between aphids on away plants and on home plants. **B** Percentage of plant-induced DE genes also appeared in the basal DE genes.

**Fig. S3.**
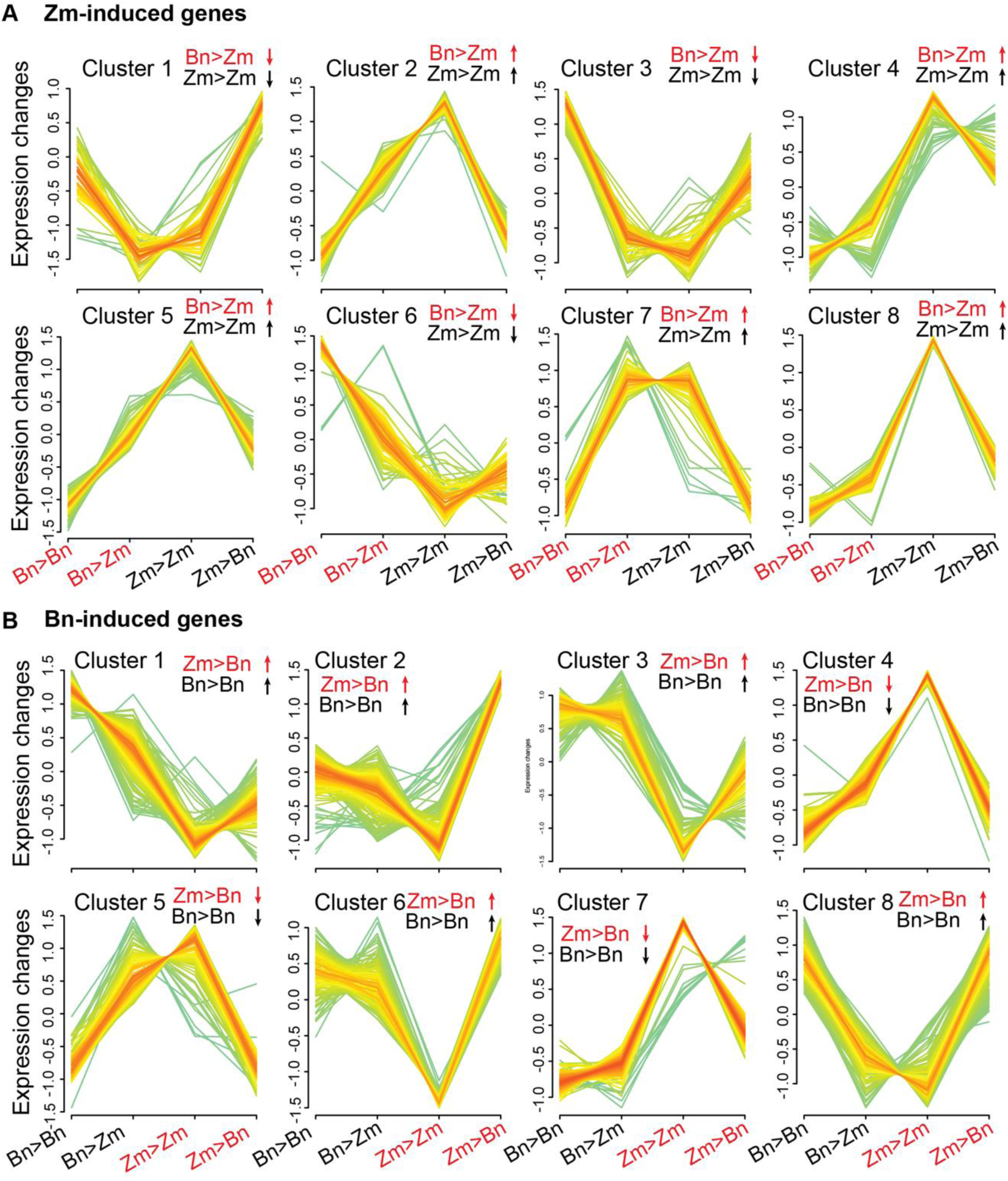
Expression pattern of plant-induced genes in GPA. **A** Expression patterns of Zm-induced genes. **B** Expression patterns of Bn-induced genes. Normalized TPM values were used to construct co-expression clusters. The colors indicate the membership belonging to the clusters. Red, yellowish, and greenish rank from high, medium, and low.

**Fig. S4.**
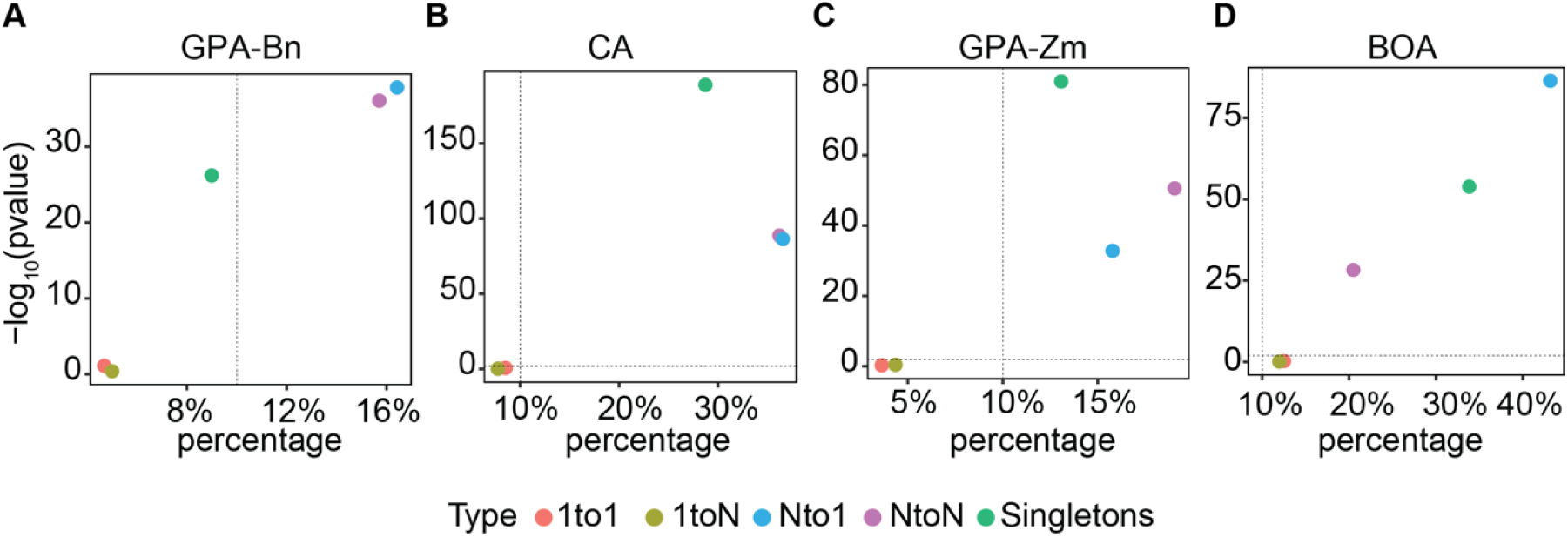
Enrichment of different types of orthologs among the randomly selected gene sets. **A**,**C** Random gene sets from GPA, **B,D** Random gene sets from CA and BOA. The X-axis is the percentage of each category in the gene sets. The y-axis is a log-transformed *p* value calculated by Fisher’s Exact test, estimating the probability of each catalog in the gene sets compared to it in all genes in a given species.

**Fig. S5.**
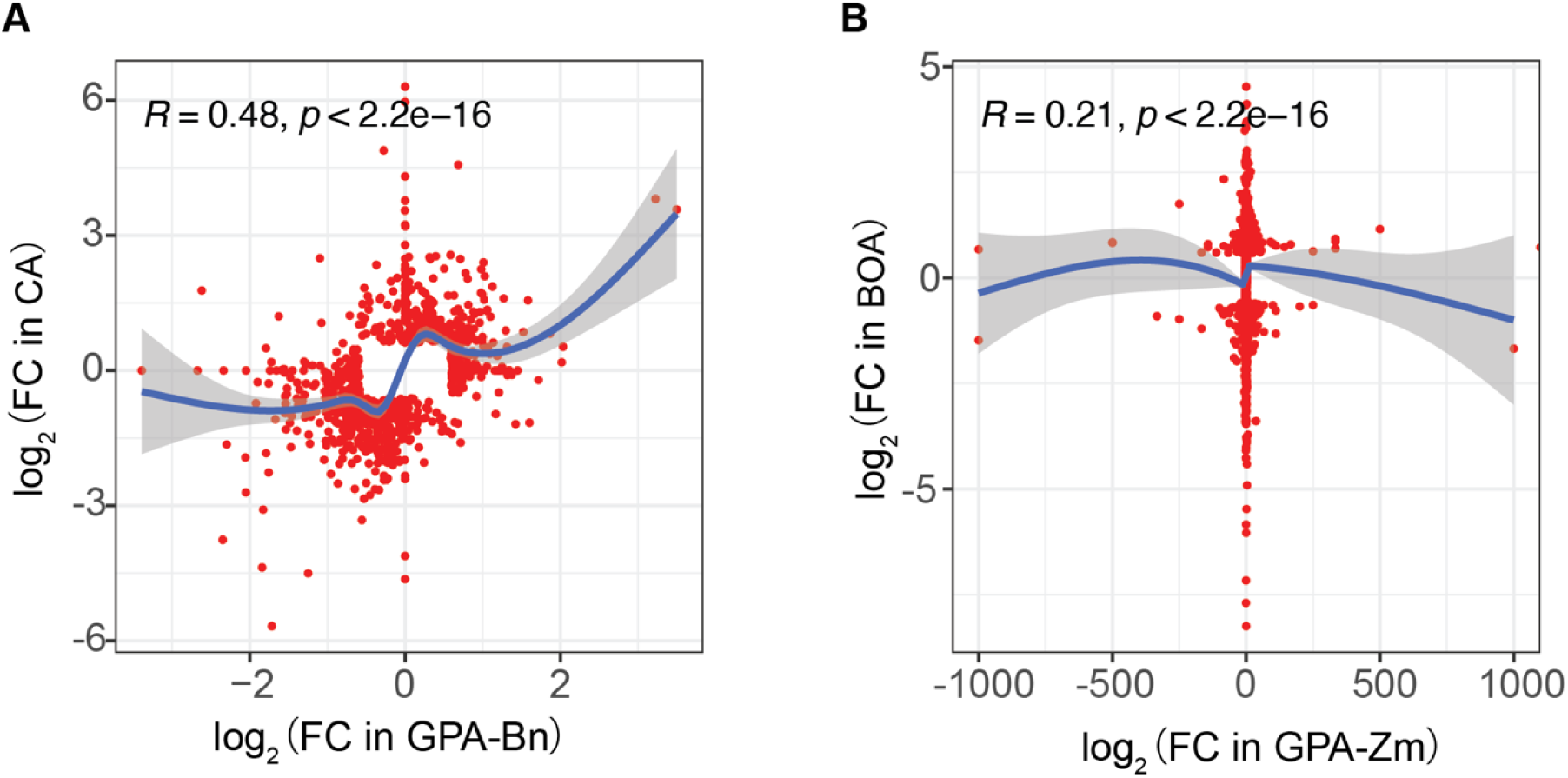
Correlation of FC of DE syntenic genes. **A** Correlation of expression changes of syntenic genes between CA and GPA that are differentially induced by Zm plants. **B** Correlation of expression changes of syntenic genes between BOA and GPA that are differentially induced by Bn plants. R is Spearman rank correlation coefficient; FC is fold-change.

**Fig. S6.**
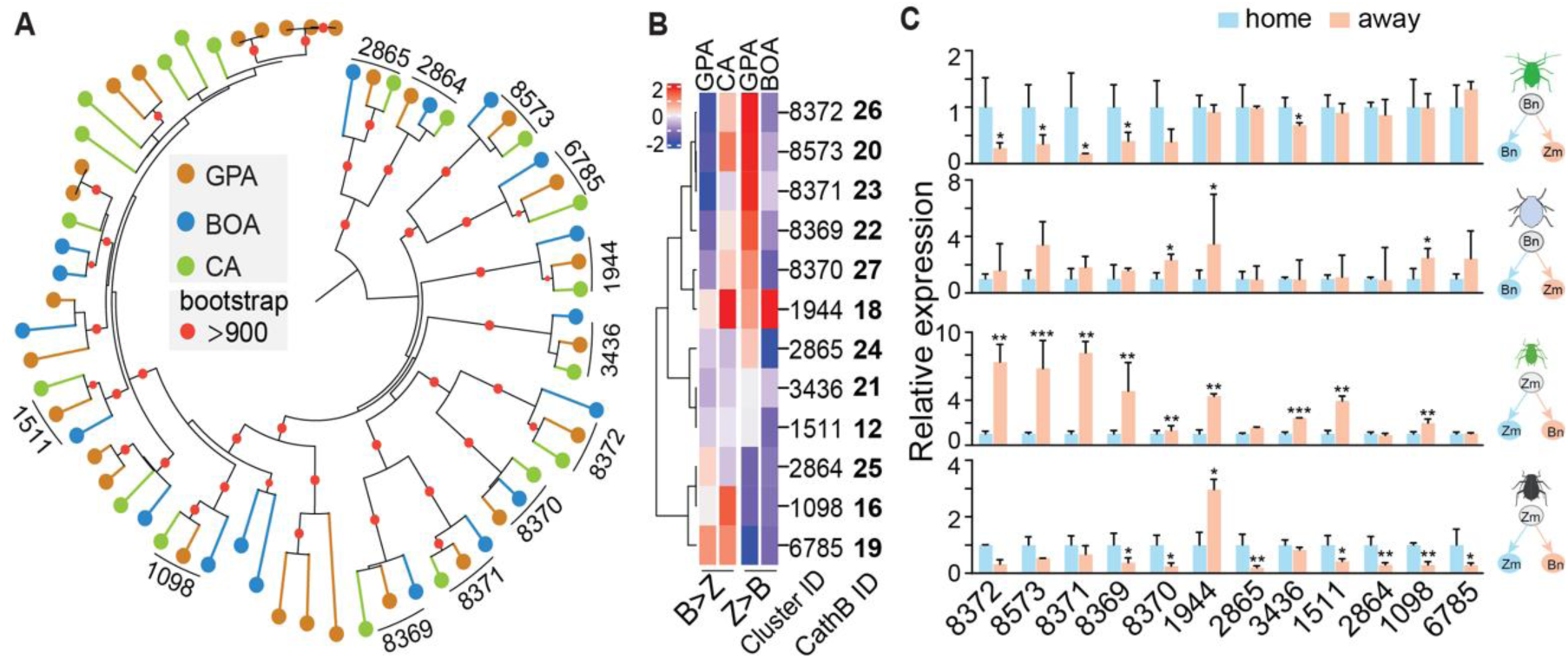
Divergent responsiveness of *Cathepsin B* genes in generalist and specialist aphids in response to plant changes. **A** Maximum likelihood phylogenetic tree of cathepsin B protein sequences in three aphid species. Syntenic copies are labeled as the cluster IDs by collinearity analysis (Dataset S3). The size of red dots in branches represents bootstrap values. **B** The column heatmaps of the fold-changes of *Cathepsin B* genes in response to plant changes. **C** Relative expression levels of *Cathepsin B* genes were quantified by qRT-PCR in three aphid species in response to plant changes. Bars represent expression values related to that of genomic housekeeping genes *EF-1α* and *RPL7* (mean± standard error) of three independent biological replicates. * *p*<0.05, ** *p*<0.01 and *** *p*<0.001 (unpaired *t* test).

**Fig. S7.**
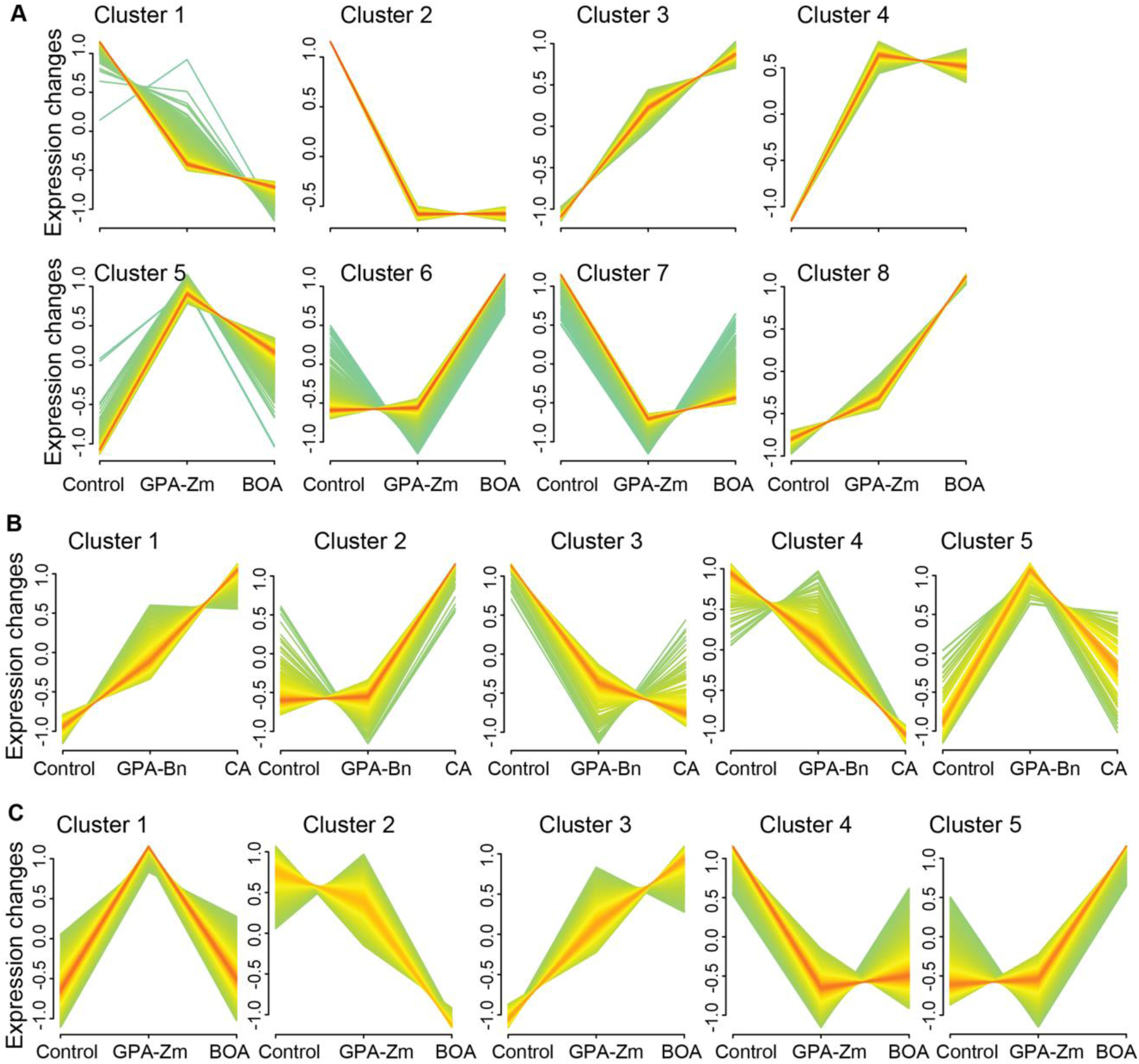
Expression patterns of plant genes induced by aphids. **A** Expression patterns of 7173 DE Bn genes. **B** Expression patterns of 997 DE Zm genes. **C** Expression patterns of 4242 DE At genes. Control is the plants weren’t treated with aphids, GPA-Zm, BOA, GPA-Bn, CA indicate the plants treated with those aphids, respectively. Experiment setups were illustrated in Fig.5. Normalized TPM values were used to construct co-expression clusters. The colors indicate the membership belonging to the clusters. Red, yellowish, and greenish rank from high, medium, and low.

**Fig. S8.**
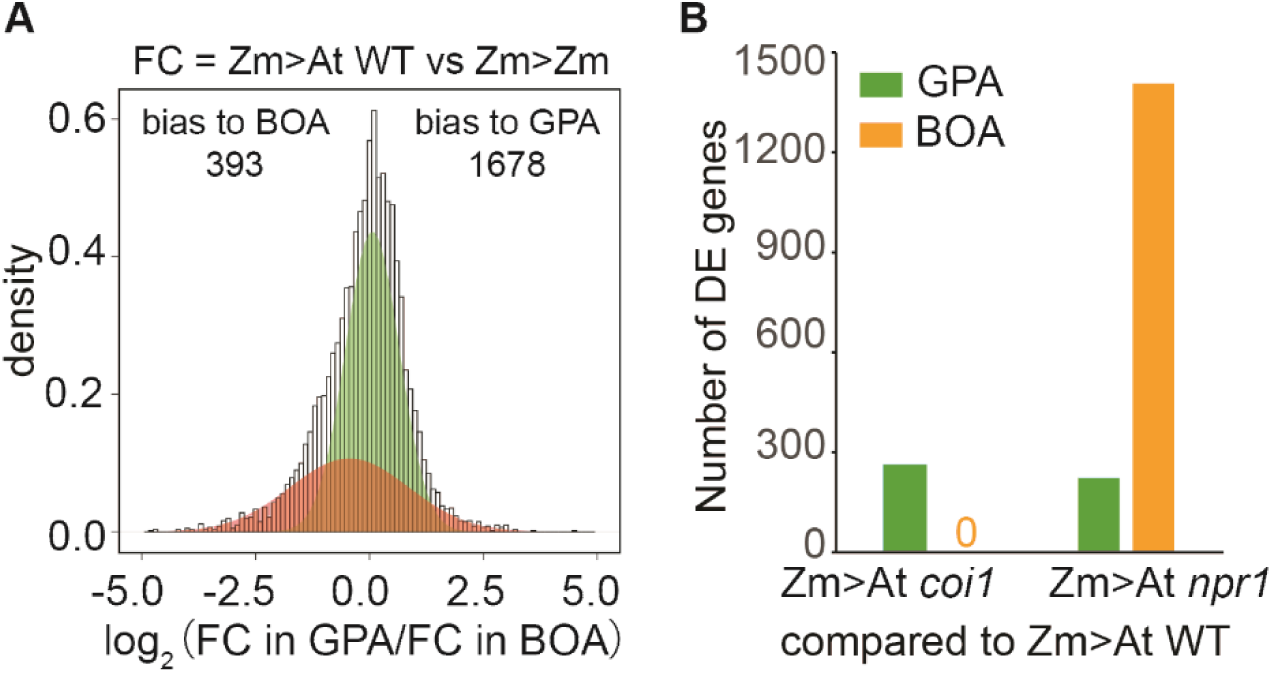
Bias-responsive genes between GPA-Zm and BOA. **A** Distribution of the fold-change ratio of syntenic genes between GPA-Zm and BOA. The light green shading represents a Gaussian component indicating unified changes of conserved genes, and the brown shading represents biased changes in one of the aphid species. **B** Number of DE genes in GPA-Zm and BOA compared aphids on Zm plants transferred to At WT and to At mutants.

### Supplementary tables

**Table S1.** Detailed information on the GO enrichment analysis for GPA-Bn genes induced by Zm plants.

| Description | p.adjust | Count | gene | total | GeneRatio |
| --- | --- | --- | --- | --- | --- |
| Hormone metabolic process | 6.06E-05 | 20 | 20 | 610 | 0.03278689 |
| Positive regulation of melanization defense response | 1.77E-04 | 7 | 7 | 610 | 0.01147541 |
| Regulation of melanization defense response | 1.96E-04 | 7 | 7 | 610 | 0.01147541 |
| Positive regulation of Toll signaling pathway | 2.64E-04 | 9 | 9 | 610 | 0.0147541 |
| Negative regulation of pri-miRNA transcription by RNA polymerase II | 4.35E-04 | 7 | 7 | 610 | 0.01147541 |

**Table S2.** Detailed information on the GO enrichment analysis for GPA-Zm genes induced by Bn plants.

| **Description** | **p.adjust** | **Count** | **gene** | **total** | **GeneRatio** |
| --- | --- | --- | --- | --- | --- |
| Cuticle development | 1.02E-07 | 45 | 45 | 692 | 0.0650289 |
| Monosaccharide metabolic process | 3.20E-05 | 47 | 47 | 692 | 0.06791908 |
| chitin-based cuticle development | 3.20E-05 | 26 | 26 | 692 | 0.03757225 |
| Secondary metabolic process | 9.84E-05 | 36 | 36 | 692 | 0.05202312 |
| Carbohydrate metabolic process | 9.84E-05 | 68 | 68 | 692 | 0.0982659 |

**Table S3.** Detailed information on the GO enrichment analysis for CA genes induced by Zm plants.

| **Description** | **p.adjust** | **Count** | **gene** | **total** | **Gene Ratio** |
| --- | --- | --- | --- | --- | --- |
| 'de novo' protein folding | 0.005563048 | 9 | 9 | 447 | 0.02013423 |
| Chitin-based cuticle development | 0.000180093 | 20 | 20 | 447 | 0.04474273 |
| Cuticle development | 0.012890953 | 25 | 25 | 447 | 0.05592841 |
| Establishment of protein localization to telomere | 0.020084221 | 6 | 6 | 447 | 0.01342282 |
| Mitotic sister chromatid segregation | 0.014646911 | 26 | 26 | 447 | 0.05816555 |

**Table S4.** Detailed information on the GO enrichment analysis for BOA genes induced by Bn plants.

| **Description** | **p.adjust** | **Count** | **gene** | **total** | **GeneRatio** |
| --- | --- | --- | --- | --- | --- |
| Cytoplasmic translation | 5.42E-22 | 72 | 72 | 1226 | 0.05872757 |
| SRP-dependent cotranslational protein targeting to membrane | 5.03E-20 | 46 | 46 | 1226 | 0.03752039 |
| Cotranslational protein targeting to membrane | 2.64E-19 | 47 | 47 | 1226 | 0.03833605 |
| Protein targeting to ER | 7.84E-19 | 49 | 49 | 1226 | 0.03996737 |
| Establishment of protein localization to endoplasmic reticulum | 1.11E-17 | 49 | 49 | 1226 | 0.03996737 |

**Table S5.**
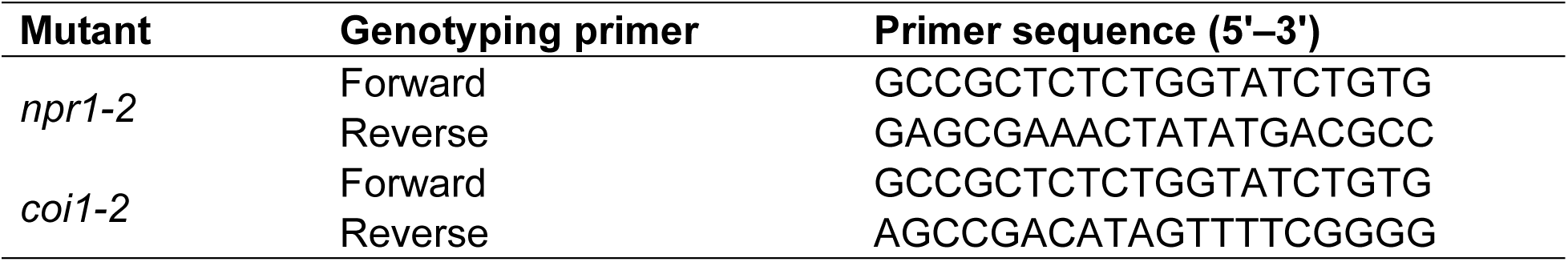
Sequences of primers used in *A*. *thaliana* genotyping experiments.

**Table S6.**
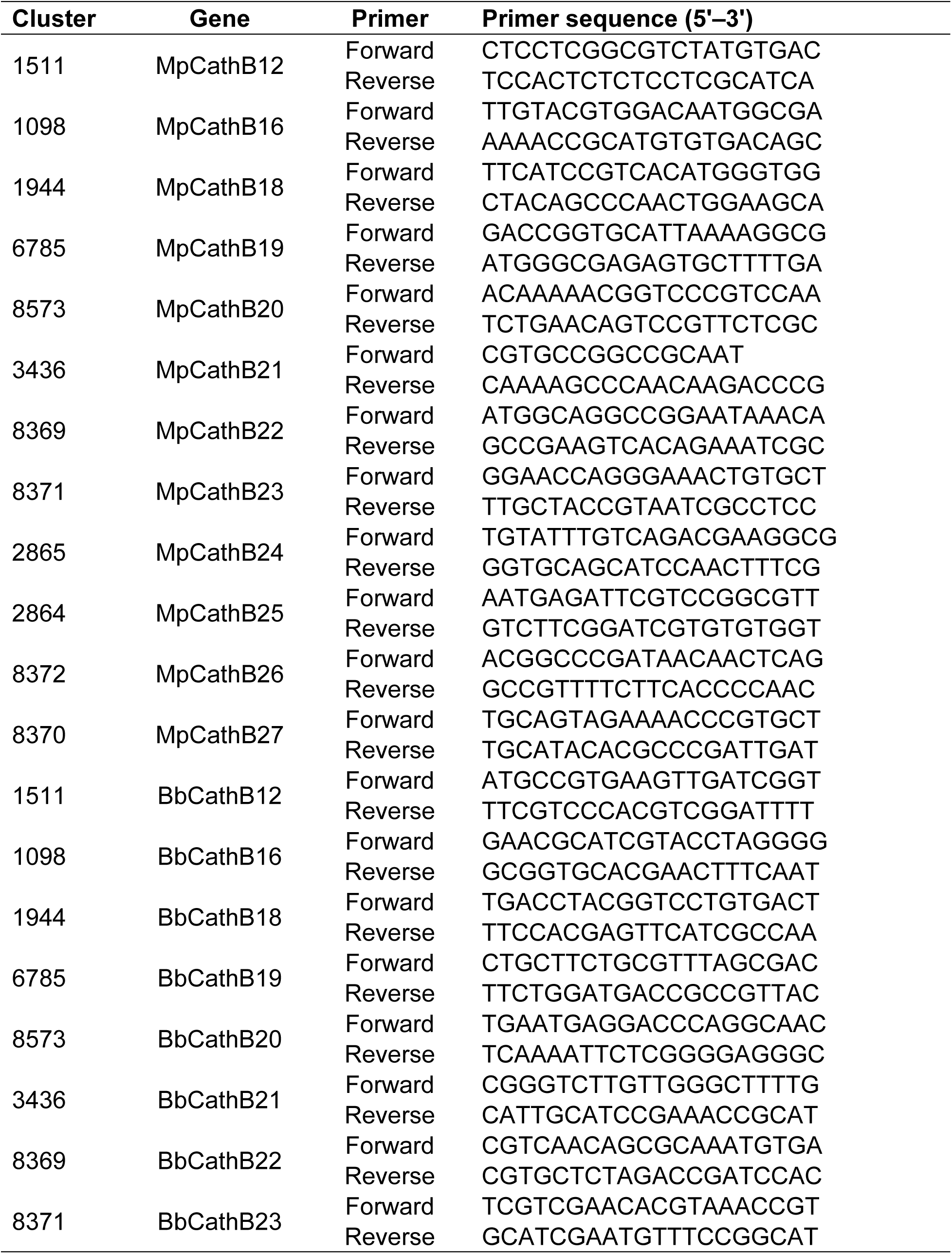
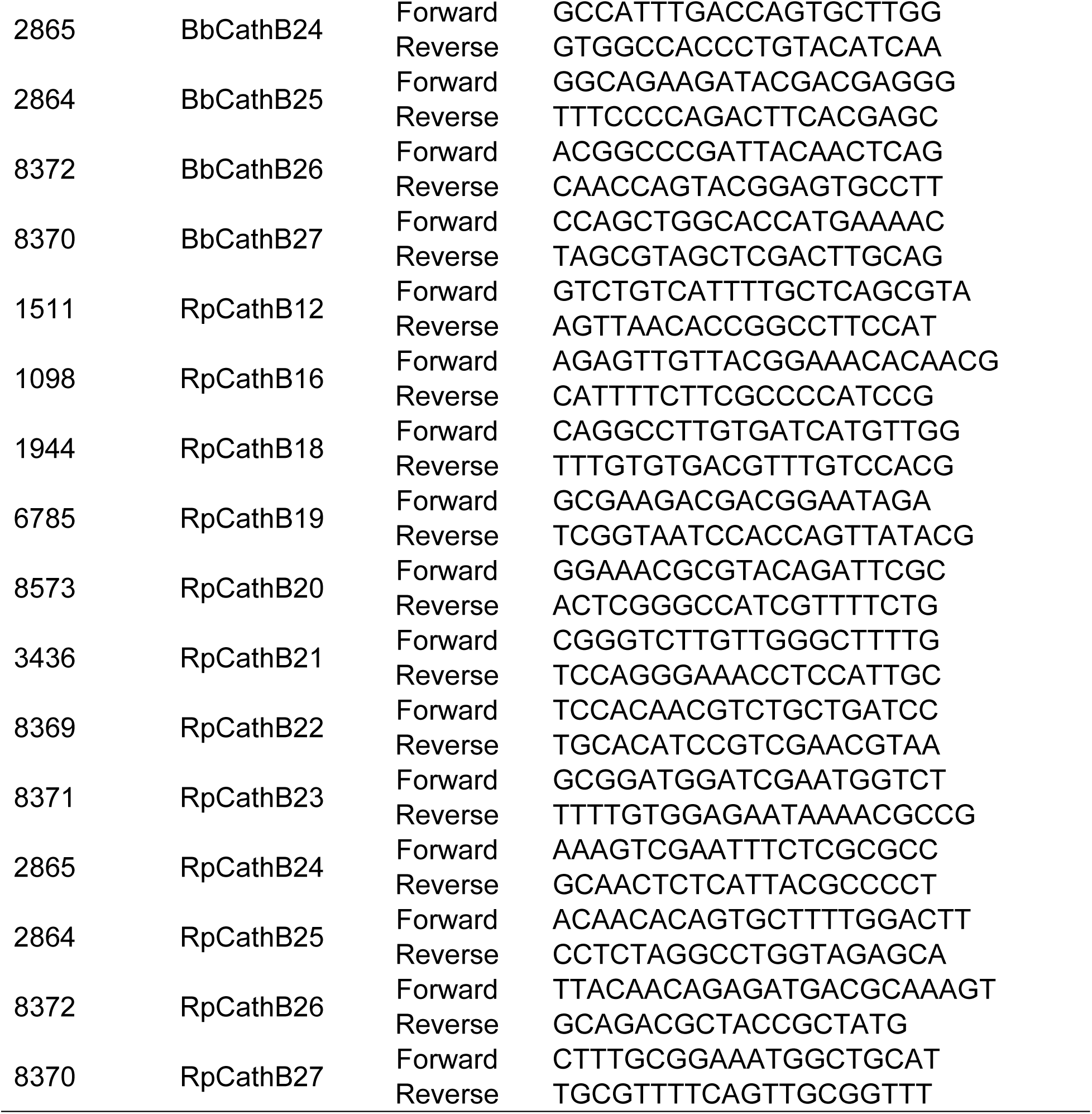
Sequences of primers used in qRT-PCR experiments.

### Legends for Datasets S1 to S10

**Dataset S1: DE genes in GPA identified in plant change experiments.**

tab 1: 3822 basal DE genes identified in GPA by the comparison between Bn>Bn and Zm>Zm.

tab 2: Zm-induced DE genes identified in GPA-Bn by the comparison between Bn>Zm and Bn>Bn.

tab 3: Bn-induced DE genes identified in GPA-Zm by the comparison between Zm>Bn and Zm>Zm.

**Dataset S2: DE genes in specialist aphids identified in response to nonhost plants.**

tab 1: 948 DE genes in CA response to nonhost plants Zm.

tab 2: 2130 DE genes in BOA response to nonhost plants Bn.

**Dataset S3: syntenic genes among three aphid species**

tab 1: 8458 syntenic genes among three aphid species.

tab 2: 1041 syntenic genes between GPA-Bn and CA that DE at least in one species.

tab 3: 2346 syntenic genes between GPA-Zm and BOA that DE at least in one species.

**Dataset S4: Biasedly responsive genes identified by the Gaussian mixture model.**

tab 1: 404 biasedly responsive genes between GPA-Bn and CA.

tab 2: 503 biasedly responsive genes between GPA-Zm and BOA.

**Dataset S5: 7309 DE Bn genes in response to aphid feeding.**

tab 1: 630 DE Bn genes in response to GPA-Zm feeding.

tab 2: 5928 DE Bn genes in response to BOA feeding.

tab 3: 4729 genes differentially expressed in Bn plants feeding by GPA-Zm compared to by BOA.

tab 4: TPM values of 7309 DE Bn genes.

tab 5: 7173 DE Bn genes were constructed into 8 co-expression clusters.

**Dataset S6: 997 DE Zm genes in response to aphid feeding.**

tab 1: 288 DE Zm genes in response to GPA-Bn feeding.

tab 2: 809 DE Zm genes in response to CA feeding.

tab 3: 204 genes differentially expressed in Bn plants feeding by GPA-Bn compared to by CA.

tab 4: TPM values of 997 DE Bn genes.

tab 5: 991 DE Zm genes were constructed into 8 co-expression clusters.

**Dataset S7: 4242 DE At genes in response to aphid feeding.**

tab 1: 1786 DE At genes in response to GPA-Zm feeding.

tab 2: 3311 DE At genes in response to BOA feeding.

tab 3: 1397 genes differentially expressed in At plants feeding by GPA-Zm compared to by BOA.

tab 4: TPM values of 4242 DE At genes.

tab 5: 4173 DE At genes were constructed into 5 co-expression clusters.

**Dataset S8: DE genes in GPA-Zm experienced plant changes from Zm to Bn, At plants.**

tab 1: DE genes in GPA-Zm that transferred from Zm to Bn plants.

tab 2: DE genes in GPA-Zm that transferred from Zm to At WTplants.

tab 3: DE genes in GPA-Zm that transferred from Zm to At *coi1* plants.

tab 4: DE genes in GPA-Zm that transferred from Zm to At *npr1* plants.

tab 5: DE genes in GPA-Zm that compared between transferring to At *coi1* and to At WT plants.

tab 5: DE genes in GPA-Zm that compared between transferring to At *npr1* and to At WT plants.

**Dataset S9: DE genes in BOA experienced plant changes from Zm to Bn, At plants.**

tab 1: DE genes in BOA that transferred from Zm to Bn plants.

tab 2: DE genes in BOA that transferred from Zm to At WT plants.

tab 3: DE genes in BOA that transferred from Zm to At *coi1* plants.

tab 4: DE genes in BOA that transferred from Zm to At *npr1* plants.

tab 5: DE genes in BOA that compared between transferring to At *coi1* and to At WT plants.

tab 6: DE genes in BOA that compared between transferring to At *npr1* and to At WT plants.

**Dataset S10: Biasedly responsive genes identified by the Gaussian mixture model.**

tab 1: 2071 biasedly responsive genes between GPA-Zm and BOA.

## Notes

### Competing Interest Statement

The authors have declared no competing interest.

